# The expression landscape and pangenome of long non-coding RNA in the fungal wheat pathogen *Zymoseptoria tritici*

**DOI:** 10.1101/2023.07.16.549241

**Authors:** Hanna M. Glad, Sabina Moser Tralamazza, Daniel Croll

## Abstract

Long non-coding RNAs (lncRNAs) are regulatory molecules interacting in a wide array of biological processes. LncRNAs in fungal pathogens can be responsive to stress and play roles in regulating growth and nutrient acquisition. Recent evidence suggests that lncRNAs may also play roles in virulence, such as regulating pathogenicity-associated enzymes and on-host reproductive cycles. Despite the importance of lncRNAs, only few model fungi have well-documented inventories of lncRNA. In this study, we apply a machine-learning based pipeline to predict high-confidence lncRNA candidates in *Zymoseptoria tritici,* an important global pathogen of wheat impacting global food production. We analyzed genomic features of lncRNAs and the most likely associated processes through analyses of expression over a host infection cycle. We find that lncRNAs are frequently expressed during early infection, before the switch to necrotrophic growth. They are mostly located in facultative heterochromatic regions, which are known to contain many genes associated with pathogenicity. Furthermore, we find that lncRNAs are frequently co-expressed with genes that may be involved in responding to host signals, such as those responses to oxidative stress. Finally, we assess pangenome features of lncRNAs using four additional reference-quality genomes. We find evidence that the repertoire of expressed lncRNAs varies substantially between individuals, even though lncRNA loci tend to be shared at the genomic level. Overall, this study provides a repertoire and putative functions of lncRNAs in *Z. tritici* enabling molecular genetics and functional analyses in an important pathogen.

**Impact statement:** Long non-coding RNAs (lncRNAs) serve distinct roles from messenger RNA. Despite not encoding proteins, lncRNAs can control important cellular processes such as growth and response to stress. In fungal pathogens, lncRNAs are particularly interesting because they can influence how pathogens infect and harm their hosts. Yet, only very few fungal pathogens have high-quality repertoires of lncRNA established. Here, we used machine learning to identify lncRNA in the major wheat pathogen *Zymoseptoria tritici.* We found that lncRNAs are highly active during the early stages of infection, before the pathogen switches to necrotrophic growth. These lncRNAs are mainly located in regions of the genome associated with pathogenicity. The repertoire of expressed lncRNAs varies substantially among individuals highlighting the potential for pathogen adaptation based on variation in lncRNAs. By expanding our knowledge of lncRNAs in important pathogen models, we enable research to comprehensively investigating their roles across fungi.

## Introduction

Long-non coding RNAs (lncRNAs) are a class of regulatory non-coding RNA (ncRNA) that lack a conserved set of features other than a length of over 200 nucleotides, and the absence of the potential to encode a functional protein [1]. As RNA molecules, lncRNAs can interact with DNA, other RNAs and proteins to regulate a wide array of molecular processes at the transcriptional, post-transcriptional, and translational levels. LncRNAs can influence the expression of genes through the recruitment of transcription factors and chromatin remodeling proteins, or through transcriptional interference [2–5]. LncRNAs may also influence the stability of a target mRNA or impact how a transcript is spliced [6]. Alternatively, lncRNAs can act as miRNA sponges or interact with ribosomes during translation [3]. The diversity of mechanisms by which lncRNAs can function is mirrored by the number of biological processes in which they intervene. In humans, many lncRNAs are differentially expressed in cancerous tissues, indicating their importance for DNA damage repair, genome stability, and the regulation of autophagy [7–9]. A notable example is MALAT1 with an unusual up-regulation in cancer cells serving as a robust biomarker for several types of cancer including in breast and lung tissue [10, 11]. LncRNAs are also important in human immunity with several loci being strongly responsive to inflammation [3, 12, 13]. Single-nucleotide polymorphisms (SNPs) in lncRNAs are also associated with various immune-related diseases such as celiac disease and atherosclerosis [14, 15]. LncRNAs tend to be more responsive to stress conditions than mRNAs [16, 17] in most organisms. In plants, lncRNAs are known to be important for the response to environmental stimuli. For example, ISP1 helps to maintain homeostasis during phosphate starvation [18, 19], and COOLAIR is an essential regulator of vernalization [19] . Naturally occurring variants of COOLAIR in populations of *Arabidopsis thaliana* require different lengths of cold exposure to de-repress flowering in early spring, and likely contribute to local adaptation [19–21]. Some lncRNAs likely contribute to the response to infection by a pathogen, through the regulation of resistance genes [22].

Interest in lncRNAs has greatly increased over the past two decades [23]. LncRNAs display higher sequence divergence and lower levels of inter-species conservation than protein-coding genes [24, 25]. Hence, lncRNAs are sometimes thought to be by-products of spurious transcription lacking any particular function [25]. Moreover, lncRNAs are typically expressed at low abundance compared to mRNAs, and often in a condition or cell-dependent manner [26], which makes their identification from sequencing data technically challenging [26, 27]. While lncRNAs can be transcribed from intergenic regions (lincRNA), lncRNAs are frequently intronic or antisense (lncNAT) to a protein-coding gene, complicating both the identification and functional validation, as knockout mutations are likely to impact not only the lncRNA but also the associated gene [28].

Most of our knowledge about the characteristics and functions of lncRNAs stems from well studied model organisms [29]. However, recently, the functions of lncRNAs across a wider range of species, including pathogenic fungi, are being explored. As in plants and animals, lncRNAs in fungi can be important regulators of growth, reproduction, and DNA damage repair [30]. In *Cryptococcus neoformans*, the switch from yeast to hyphal growth before sexual reproduction is an important component of virulence, and the upstream lncRNA RZE1 regulates the main gene (ZnF2) responsible for this switch in morphology [31]. In *Candida auris*, the deactivation of the lncRNA DINOR results in higher levels of DNA damage and constitutive filamentous growth [32]. In *Cryptosporidium parvuum,* 86% of all predicted lncRNA candidates are differentially expressed between sexual and asexual phases [33]. Fungal lncRNAs are also highly responsive to environmental conditions, particularly in response to stress [30, 34]. Predicted lncRNAs in the insect pathogen *Metarhizium robertsii* show high levels of differential expression during heat stress, with many predicted targets being directly implicated in responding to the signal [34]. In *Ustilaginoidea virens*, fine-tuned transcription of the lncRNA UvMFS is required for growth under stress [35]. In *Candida*, lncRNAs were shown to be differentially expressed during the infection of epithelial cells [32]. Several lncRNAs with direct effects on virulence were discovered recently, including in *Trichoderma reesei,* where natural variants in the lncRNA HAX1 influence cellulase production [36]. In *C. neoformans,* lncRNAs were found in extracellular vesicles containing virulence factors known to modulate the host immune response [37]. In *Verticilium dahliae*, three lncRNAs were found to regulate the expression of cell-wall degrading enzymes, with mutants resulting in decreased virulence on cotton [38]. In *Fusarium graminearum*, the lncRNA lncRsp1 influences virulence on wheat by regulating *Fgsp1*, which is required for normal ascospore discharge [39]. These examples illustrate the multitude of roles lncRNAs can play in pathogenicity-associated processes of filamentous fungi.

*Zymoseptoria tritici* is a filamentous ascomycete and the causal agent of Septoria blotch, one of the most important crop diseases worldwide [40]. *Z. tritici* populations show high levels of genetic diversity even at small geographical scales [41, 42], coupled with significant variability in gene expression between isolates [43, 44]. The genome contains a high number of transposable elements (TEs) [45, 46] which may provide frequent opportunities for the formation of functional lncRNAs [47]. During infection on wheat, *Z. tritici* undergoes a switch from biotrophic to necrotrophic lifestyle, and a vast transcriptional reprogramming is required for the necessary metabolic changes [43, 44]. Significant morphological changes have also been observed during growth in stressful conditions [48]. Epigenetic control of TE-rich, accessory regions likely contributes to high levels of expression variation of infection-related genes [49] An increase in TE expression and an enrichment of small RNAs (sRNAs) originating from accessory chromosomes was observed during growth in nutrient-poor conditions, indicating a potential role of these regions in the response to stress [50]. Despite a clear transition in the sRNA transcriptome under stressful conditions, no direct role in host colonization has been demonstrated for sRNAs, in contrast to several other fungal plant pathosystems [51] . Recently, an improved genome annotation based on long-read transcript sequencing was established [52] reporting the production of several lncRNAs during growth *in-vitro.* The study further demonstrated that lncRNAs were differentially expressed between *in-vitro* and *in-planta* conditions, and that some lncRNAs showed interesting expression correlation patterns with nearby genes during infection [52].

In this study, we identify and assess high-confidence candidates to reveal the landscape of lncRNAs in the *Z. tritici* genome, identify biological processes associated with lncRNA functions and assess expression variation over an infection life cycle. Finally, we assess pangenome features of lncRNA loci to define conserved and variable elements of the genomic and transcriptomic landscape of lncRNA.

## Methods

### LncRNA candidate identification

Raw RNAseq data from the *Z. tritici* reference isolate Zt_09 (IPO323ΔChr18) produced at four stages of the infection cycle *in planta* was downloaded from NCBI (accession PRJNA415716) [44]. Using the reference genome annotation for IPO323 [53], we used the tool PINC [54] to predict lncRNA candidates from transcriptomic data. We set the weight to 0.7 on the Youden’s index instead of the default 0.5 in order to decrease false-positive rates [54]. We included consensus sequences of annotated TEs [45]in the file containing known protein-coding mRNAs, in order to reduce the number of candidates originating from degraded TE insertions. We retained only the longest transcript at each locus. The GFF annotations for predicted lncRNAs was compared using gffcompare (v0.11.2) [55] to the reference annotation for IPO323.

### Differential expression

RNAseq reads were aligned to the reference genome IPO323 using Hisat2 [56] with default parameters and exon-level counts were quantified with featureCounts (v2.0.1) [57] using default parameters. FeatureCounts was run twice; once using the GTF file provided by the output of PINC and once using the reference annotation, in order to more accurately quantify expression of reference genes that were missing in the stringTie (v2.2.1) [58] assembly performed within PINC. Using the *edgeR* package (v3.17) [59] in R (v4.3.1) [60], lncRNAs and genes were tested for differential expression between the two earliest time-points and the two latest. The FDR cut-off was set to 5%. In order to compare expression between isolates, reads were aligned to each respective reference genome (*i.e.,* 1A5, 1E4, 3D1, and 3D7) [61, 62] using Hisat2 with default parameters. Mapped reads were quantified with featureCounts using the GTF file containing the PINC predictions.

### Clustering

Trimmed Mean of *M*-values (TMMs) of genes and lncRNAs were assessed by edgeR and scaled to achieve comparable variance across the four time-points. Values of both genes and lncRNAs were clustered according to their trajectories across time-points using fuzzy c-means clustering using the *e1071* (v1.7-13) [63] package in R. Helper functions [64] were used to find the optimal hyper-parameters. Within-sum-of squares indicated that the optimal number of clusters was 7. After clustering was performed, clusters were qualitatively grouped into three groups based on peak expression: early expression peak, late expression peak, and bi-modal expression peak (showing peaks at both early and late infection with reduced transcription at intermediate time-points).

### Target prediction and correlation

All transcripts within 50kb of a lncRNA were extracted from the reference genome using bedtools (v2.30.0) [65]. lncTAR [66] was used to predict potential interactions between lncRNA/mRNA pairs in this window. Pairs were considered to have a potential interaction if the ndG was lower than the default -0.1. For each lncRNA, the expression of each transcript within a 50-kb window was correlated to the expression of the lncRNA using the scaled TMM values. Differences between interacting and non-interacting pairs were analyzed in R. For selected lncRNAs, RNAfold was used to predict the secondary structure with default parameters. Functional domains encoded by the interacting genes were analyzed from previous annotations [53]. Functional enrichments were performed using the *Gostats* package (v3.17) [67] in R. Biosynthetic gene clusters of the reference genome *Z. tritici* (IPO323) were predicted using antiSMASH v.7.0 [68].

### Chromatin immunoprecipitation (Ch-IP) sequencing

We compared histone modification H3K27me3 and H3K4me2 profiles of the reference genome of *Z. tritici* (IPO323) cultured in carbon limited medium (minimal medium) and carbon rich medium (YSB). We performed culturing for ChIP-seq analyses of *Z. tritici* based on growth in Vogel’s Medium N (minimal medium) until hyphae formation for 8 days at 18°C. The ChiP-seq library was prepared for sequencing and analyzed using a NovaSeq 6000 in paired-end mode with a read length of 150 bp. ChIP-seq data of *Z. tritici* grown in YSB medium was retrieved from the NCBI SRA database (accession number SRP059394; ChIP-seq raw reads were trimmed using Trimmomatic v.0.32 [69] with parameters ILLUMINACLIP:2:30:10 LEADING:3 TRAILING:3 SLIDINGWINDOW:4:15 MINLEN:36. and mapped to the reference genome IPO323 with Bowtie2 v.2.4 [70] --very-sensitive-local parameter. Duplicated reads were tagged with the GATK Picard MarkDuplicates function v.4.2.4.1 . Peak calling was performed with Epic2 v.0.0.52 [71] setting bin-size 1000 –mapq 5.

### Pangenome analyses

Fasta sequences of filtered lncRNA candidates were extracted from the transcriptome assembly provided by PINC using bedtools, and were aligned to the genome sequence of 18 alternate reference isolates using exonerate (v2.70.2) [72] (model est2genome with a maximum intron length of 300). Transcripts returning a significant match to a particular genome were considered to be present. To estimate the number of additional lncRNAs that could be present on a population scale, the steps outlined in candidate identification were for 4 other isolates (1A5, 1E4, 3D1, and 3D7). The resulting transcript sequences were clustered with cd-hit (v4.8.1) [73] using relaxed parameters (sequence identity of ≥80%, minimum alignment coverage ≥75%). Transcripts belonging to the same cluster were considered as the same lncRNA locus regardless of genomic location and context. The accumulation curve was constructed using the clusters of transcripts using the *vegan* library (v2.6-4) [74] in R. Correlation plots were created using the corrplot package in R [75]. All additional statistical tests were also performed in R and all plots were created using the ggplot2 package v3.4.2 [76].

## Results

### Identification and characterization of lncRNA candidates across the genome

We predicted lncRNA loci based on transcriptomic datasets collected across multiple environments. RNAseq data from the *Z. tritici* reference isolate Zt09 generated at four stages of the infection cycle on wheat plants was mapped to the reference genome IPO323 [44]. Zt09 is a derivative (IPO323ΔChr18) of the reference strain differing only by the deletion of the accessory chromosome 18 ^41^. We used the prediction pipeline PINC [54] to predict lncRNA candidates, integrating the consensus sequences of all known TEs as well as all annotated genes as known mRNA sequences. This reduced the number of predicted lncRNAs originating from TEs. We obtained a total of 120 putative lncRNA candidates originating from 108 distinct loci. Of the 12 loci predicted to produce multiple lncRNAs, six were predicted to produce more than one transcript of exactly the same length. While lncRNA isoforms are known to exist in other organisms [77], isoforms are not expected to show the same transcription start and stop site, as well as being of exactly the same length. Furthermore, RNA-isoform-level downstream analysis requires care and cannot be fully resolved using single-stranded RNA-seq data [78]. In order to facilitate downstream analysis and avoid false calls originating from transcript assembly, we filtered the original lncRNA candidates to retain only the longest transcript at each locus. This resulted in a total of 91 predicted lncRNAs [Supplementary table 1].

A revised IPO323 genome annotation based on culture medium Iso-Seq libraries predicted 51 lncRNAs [52]. Only two of our 91 lncRNA (2.2%) candidates overlap with these predictions. Low congruence between these two pipelines may be a result of several factors: the authors excluded all transcripts under <1kb and containing an ORF longer than 300bp [52], which differed from our pipeline that uses two machine-learning models to assess coding potential in relation to known coding sequences instead of filtering by ORF length [52]. Furthermore, expressed lncRNAs may be significantly different between *in vitro* and *in-planta* conditions. Although the vast majority of our candidates were not predicted to be lncRNA in this new annotation, our candidates were generally supported by the Iso-Seq data. Despite being only from culture medium condition, a total of 62.8% of our candidates have at least one fully covering read, and 78.26% show non-zero coverage over more than 75% of their length [Supplementary figure 1].

LncRNA loci were found on most chromosomes with the exception of chromosomes 10, 16, 17, or 20 [Supplementary table 1]. Chromosomes carried 1-13 lncRNAs with the highest number found on chromosome 7. The large majority (91.1%) of lncRNAs were located on core chromosomes (excluding chromosome 10). The maximum number of lncRNAs found on an accessory chromosome was 4 (chromosome 19) [Supplementary figure 2]. The distribution of lncRNAs along the chromosomes showed no apparent associations with chromosomal features with loci being discovered both in non-telomeric and sub-telomeric regions [Figure 1A]. The density of lncRNAs was largely dependent on the total size of the chromosome, except for chromosome 7, which not only showed the highest number of lncRNAs overall, but also the second highest density of lncRNAs on the core chromosomes despite the relatively large size [Figure 1B]. Out of the 91 lncRNAs, 87 were classified as lincRNAs while only 3 were lncNATs [Figure 1C]. Compared to genes, lncRNA loci encode in general fewer exons [Figure 1D] (mostly 1-2 exons) and transcripts were slightly shorter on average [Figure 1E], which is consistent with knowledge about lncRNAs in other organisms [23].

**Figure 1:**
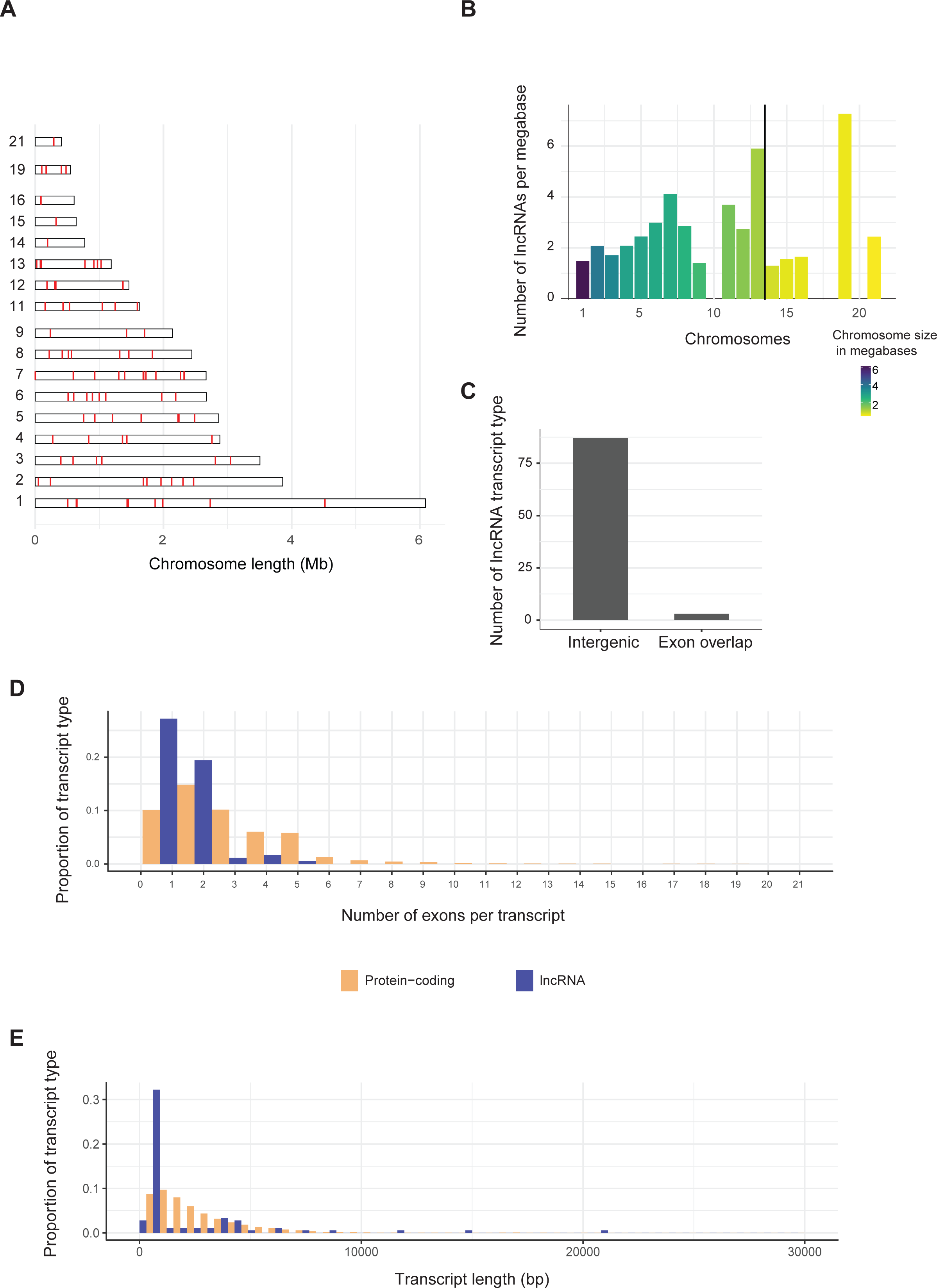
A) Schematic representation of the distribution of lncRNA along chromosomes. Each red line represents a predicted lncRNA. B) Density of lncRNAs on each chromosome. C) Bar chart of the distribution of lncRNAs into different types based on genomic context. “Intergenic” indicates that the lncRNA originates from a locus that has no overlap with a known coding sequence. “Exon overlap” indicates that the lncRNA shares at least one exon with a known mRNA. D) Histogram comparing the the number of exons per transcript between lncRNA and mRNA. E: Histogram comparing the transcript length in basepairs between lncRNA and mRNA.

### Genomic niches and expression dynamics of lncRNA loci

In order to understand the genomic context of the predicted lncRNAs compared to mRNAs encoded by genes, we assessed distances of lncRNA loci to nearest neighboring genes. Compared to genes, lncRNAs were not at a significantly different distance to the nearest gene in any individual orientation (sense upstream/downstream, antisense upstream/downstream) [Figure 2A]. Moreover, there was no significant difference in the distance to their nearest neighbor regardless of the direction [Figure 2B]. Next, we assessed if lncRNAs were more likely found in a particular orientation relative to their nearest neighboring gene [Figure 2C]. We found no significant differences between lncRNAs and mRNAs (Chi-squared test; Chi-squared = 2.8395, df = 3, *p*-value = 0.417) indicating that, lncRNAs are found in similar genetic contexts as protein-coding genes.

**Figure 2:**
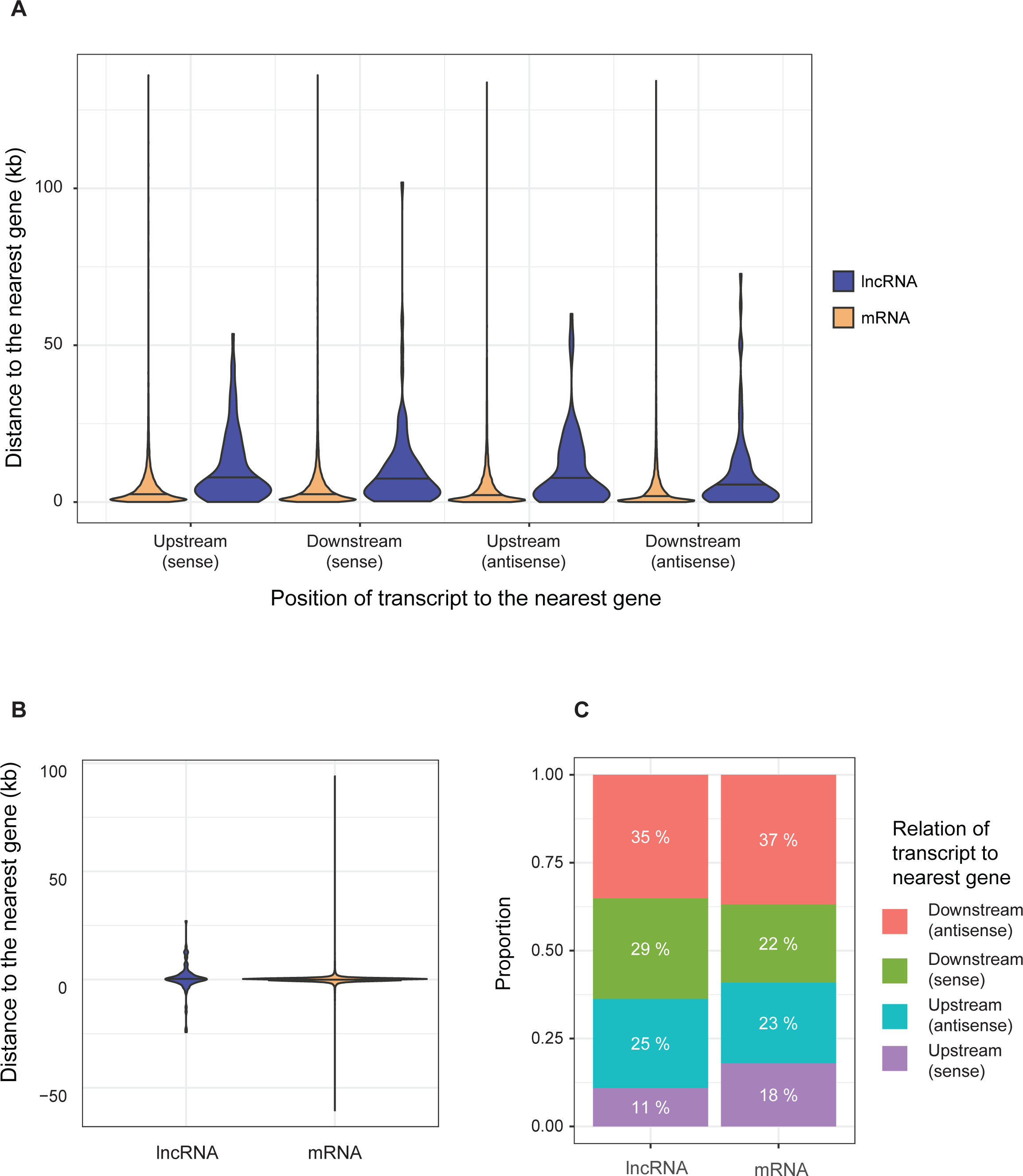
A) Violin-plot comparing the distribution of distances to the nearest neighboring gene in each orientation for lncRNA and mRNA. Upstream and downstream refer to the position of the lncRNA or mRNA relative to the neighbor. Sense and antisense refer to whether or not the lncRNA or mRNA is transcribed from the same strand as the neighbor. B) Violin plot comparing the distribution of the distance to the nearest neighboring gene, regardless of orientation. C) Comparison of the orientation of the nearest neighboring gene between lncRNAs and mRNAs.

LncRNAs are known to be expressed at lower levels than mRNAs [23]. To test this in *Z. tritici*, we assessed TMMs by infection time-point for both lncRNAs and mRNAs. As expected, lncRNAs were significantly less expressed at all four time points capturing the infection lifestyle transitions with an average effect size of -2.6 compared to protein-coding genes [Figure 3A]. LncRNA and mRNA expression differences were assessed in four additional strains with comparable RNA-seq datasets. LncRNAs were expressed at consistently lower levels compared to mRNA [Supplementary figure 3]. Intraspecific variation in lncRNA expression is known to be higher than for protein-coding genes across kingdoms [16, 79, 80]. To test for such differences, we compared lncRNA and mRNA expression variation between four individual strains of *Z. tritici.* We used the coefficient of variation for each transcript between strains at each time-point to define expression variation. We removed all transcripts with <10 counts per million (CPM) in one or more of the three biological replicates for each strain and time point to reduce noise originating from lowly expressed loci. LncRNAs show a higher variability in expression than mRNAs at all time-points, with the strongest effect at 28 days post infection (dpi) [Figure 3B].

**Figure 3:**
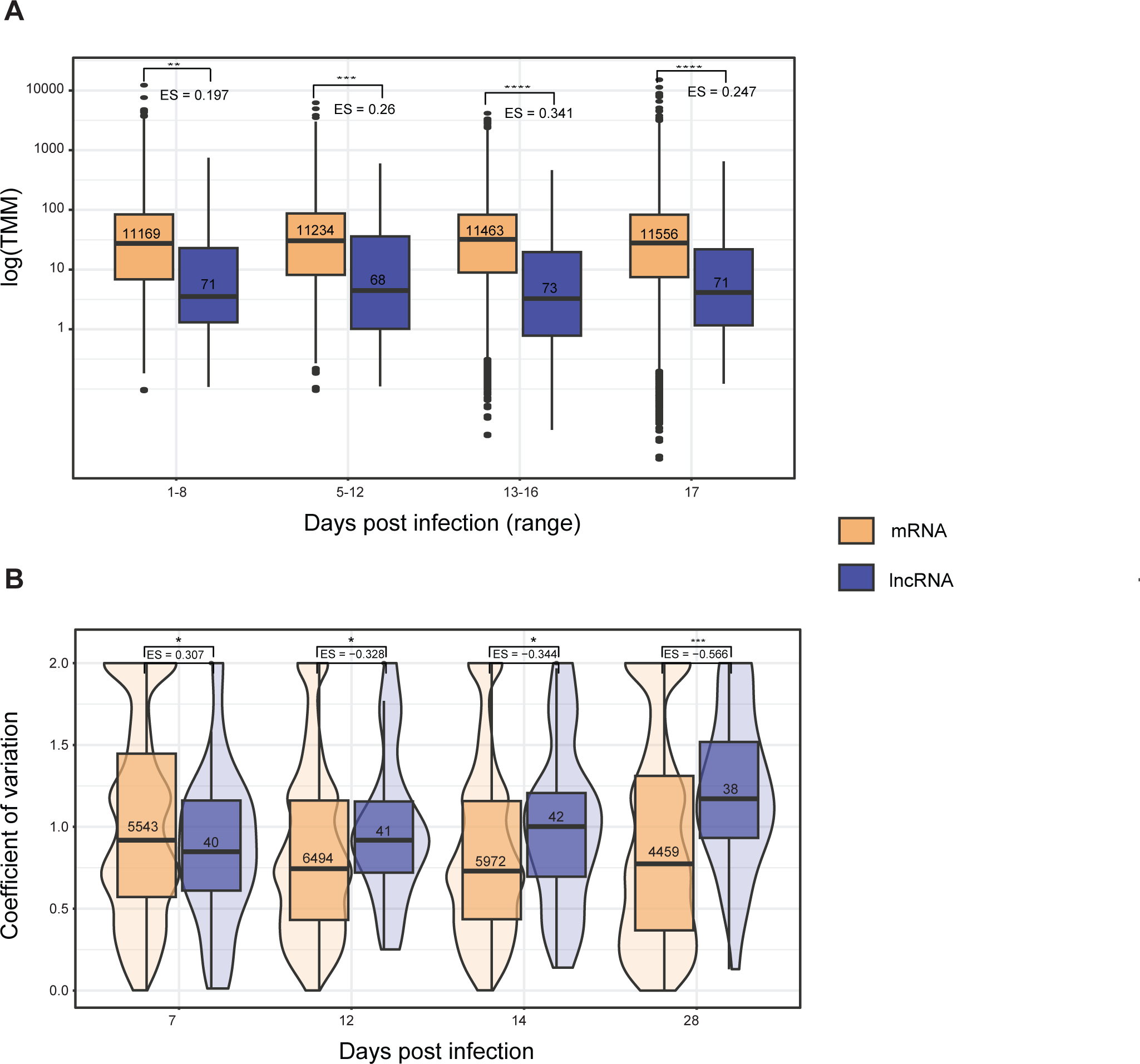
A) Distribution of log TMM values between lncRNA and mRNA. Stars show the significance levels of Welch two-sided *t-*tests comparing the mean of each transcript type at each time point. Values underneath the brackets (denoted by “ES=”) show the effect size relative to mRNA expression. Values inside the boxes show the sample size of each group. B) Violin plot comparing the distribution of the coefficient of variation (CV) for each transcript, calculated by comparing centered and scaled TMM values between four isolates for mRNAs (red) and lncRNAs (blue) at each time-point. Stars show the significance levels of Welch two-sided *t-*tests comparing the mean CV between the two transcript types. Values denoted by “ES=” represent the effect size relative to the mean mRNA CV at each time-point.

### Potential regulatory targets of lncRNA

Regulatory effects of most known lncRNAs are acting in *cis* with a minority acting in *trans* [81]. However, targets in *cis* are also computationally less demanding to detect, given the smaller search space for associations across the genome. To identify the range of potential *cis* targets of the identified lncRNAs, we extracted all annotated genes within a 50 kb window around each lncRNA-encoding locus. Given the average distance between genes of ∼1 kb, the window typically contains dozens of genes. Compared to the rest of the genome, genes within the window are enriched for molecular functions involving catalytic and enzymatic activity as well as binding functions (most significantly ATP and lipid binding) [Supplementary figure 4]. In terms of biological processes, genes near lncRNAs are enriched in functions related to protein metabolism and modification processes [Supplementary figure 5].

Proximity to genes is insufficient to ascertain *cis-*acting lncRNA functions, hence we assessed whether expressed genes and potential lncRNA regulators were positively or negatively correlated within a range of 50 kb. Overall, no trend in correlation coefficients was found based on the distance of lncRNAs to mRNAs, and no significant differences were found in lncRNA-mRNA pairs within 50 kb compared to the background, indicating that proximity alone does not drive lncRNA-mRNA co-expression [Figure 4A; Supplementary table 2]. LncRNAs may modulate the expression of nearby genes by interacting with an mRNA to form RNA-RNA duplexes [82, 83], and the potential for two RNAs to interact can be predicted by free-energy minimization [84]. Using the software lncTAR [66], we analyzed the interaction between lncRNA and mRNAs in the same 50 kb windows. We found that 43.3% of lncRNAs had no predicted RNA interaction, while 45.6% of lncRNAs had predicted interactions with 1-5 mRNAs. The maximum number of predicted lncRNA-mRNA interactions in a 50-kb window was 23 [Figure 4B]. We compared the correlation of expression values between interacting and non-interacting pairs [Figure 4C]. We found that pairs of lncRNAs and mRNAs that could interact were significantly more anti-correlated with each other than pairs that were not predicted to interact as long as the lncRNA-mRNA pair was located within 1 kb (effect size 1.085) [Figure 4D]. At longer distances, the potential for the transcripts to interact with each other was not significantly associated with expression correlation [Figure 4D].

**Figure 4:**
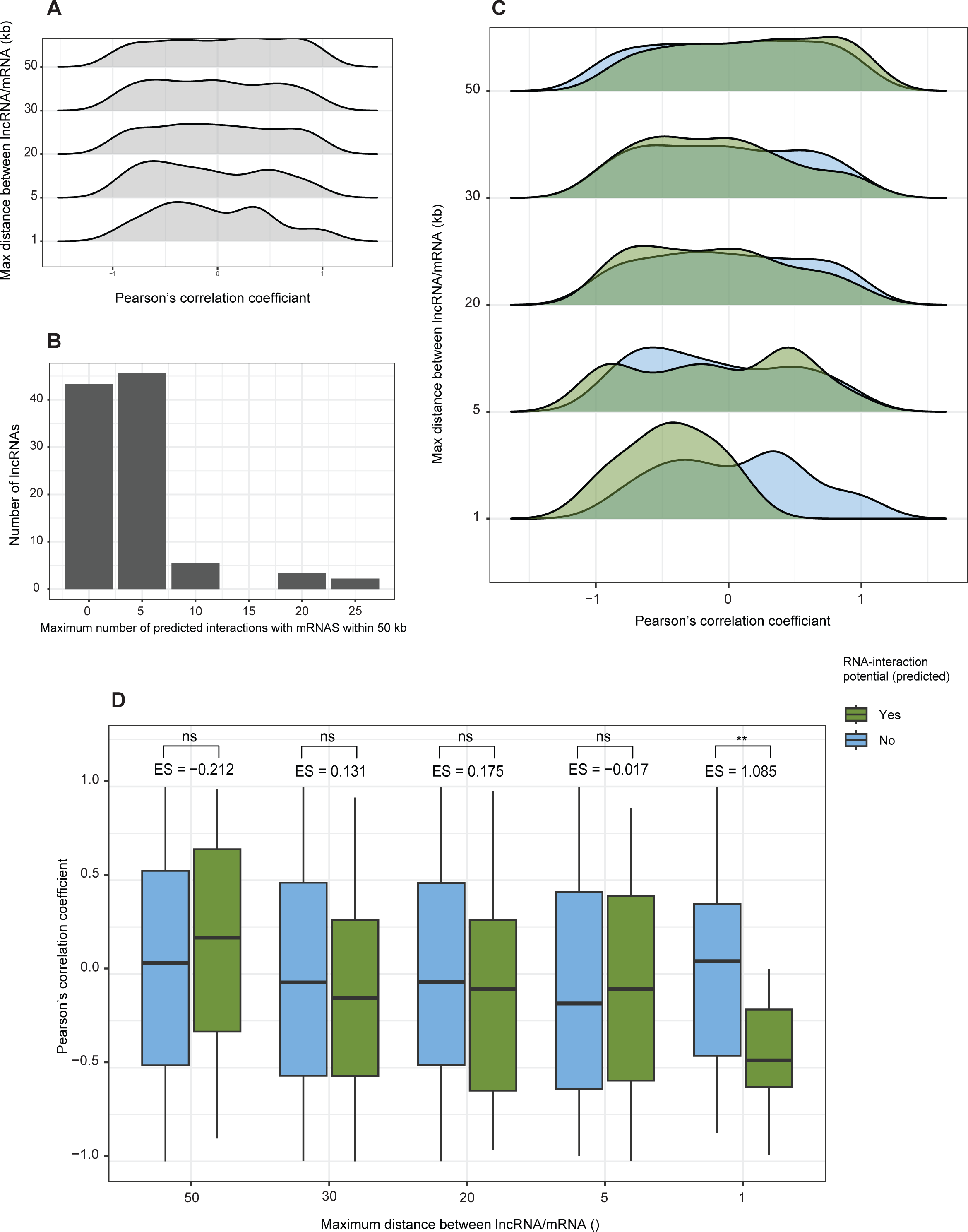
A) Density plots comparing the distribution of correlation coefficients (Pearson’s *r*) between lncRNAs and mRNAs, based on the distance between the pairs. The maximum distance between the pairs is represented on the y-axis, with the minimum being the boundary of the next smaller distance category. For example, the distribution shown at 50 kb represents all lncRNA-mRNA pairs between 30-50 kb. B) Total number of significant lncRNA-mRNA interactions per lncRNA, predicted by free energy minimization. C) Density plots comparing the distribution of correlation coefficients (Pearson’s *r*) based on the distance between the pair and their potential to interact. Pairs with no predicted interaction are shown in blue, while pairs with a predicted interaction are shown in green. D) Distribution of correlation coefficients (Pearson’s *r)* based on the distance between the pair and their potential to interact. Pairs with no predicted interaction are shown in blue, while pairs with a predicted interaction are shown in green. Stars represent the significance levels of Welch two-sided *t-*tests comparing the mean correlation coefficient of interacting and non-interacting pairs at each distance. Values denoted by “ES=” show the effect size relative to the correlation coefficients of non-interacting pairs.

To investigate gene functions potentially regulated by lncRNAs, we performed an enrichment analysis of all predicted mRNA targets at a maximum distance of 5 kb and showing an absolute correlation coefficient >0.5 with the interacting lncRNA against the genomic background. Most enriched biological processes were related to oxygen stress and detoxification [Supplementary figure 6]. The strongest enriched molecular function was antioxidant activity [Supplementary figure 7]. Among all potential lncRNA targets, regardless of distance or correlation, enriched biological pathways include regulatory processes involved in homeostasis, as well as translation, ion transportation, and metabolism [Supplementary figure 8]. Metal-ion transmembrane transporter activity is the most strongly enriched molecular function among all the potential mRNA targets compared to the genome [Supplementary figure 9], and it is also enriched when compared to non-target genes within the same 50 kb interval [Supplementary figure 10].

### Differential expression of lncRNAs during plant infection

The infection cycle of *Z. tritici* on wheat includes four distinct morphological stages characterized by the up-regulation of particular gene functions and pathways [43, 44, 85]. *Z. tritici* isolates are highly diverse both genetically and transcriptionally, and as a result the timing of each stage varies significantly between isolates [43, 44]. For simplicity, we refer here to the timing of each stage according to the reference isolate Zt09. At the earliest stages (1-8 days post infection, dpi), spores germinate on leaves and hyphae enter through the stomata. From 5-12 dpi, the pathogen colonizes the mesophyll concluding the biotrophic stages. During biotrophic growth, genes involved in lipid catabolism are up-regulated, indicating that the pathogen is relying on internal energy storage [85]. The highest number of predicted effectors are up-regulated at the 5-12 dpi stage [85]. From 13-16 dpi, the pathogen forms pycnidia as it begins to acquire nutrients from the host and enters the necrotrophic stage, characterized by the up-regulation of cell-wall degrading enzymes, transmembrane transporters, and genes involved in secondary metabolite production. Beyond ∼17 dpi, the pathogen can produce pycnidia inside the stomatal cavity [44].

We classified lncRNA transcription peaks according to the specific infection stages, using fuzzy c-means clustering on all gene and lncRNA expression trajectories across all time points [Figure 5A]. Both genes and lncRNAs were found in all seven clusters. Clusters with similar expression profiles were grouped to obtain three groups of clusters with expression peaks during early infection, late infection, or with high expression at both early and late time points [Figure 5B]. LncRNAs were more likely to show peak expression during early infection compared to genes (Chi-squared test; *p*-value = 0.01015) [Figure 5C]. Clusters 4 and 5 contained the two highest numbers of lncRNAs and both showed peak expression during early infection. Cluster 5 includes the highest number of lncRNAs, and is enriched in genes encoding nucleic-acid binding domains, involved in catalytic activity, and oxidoreductase / peroxidase / antioxidant activity [Supplementary figure 11], which have been shown to be important for overcoming the host defense response [86, 87]. Secreted peroxidases have been shown to be important pathogenicity factors required for symptom formation on wheat by *Z. tritici* [88]. Cluster 4 shows an enrichment of functions related to serine peptidases, sulfur transmembrane transporters, and vitamin B6/pyridoxine binding [Supplementary figure 12]. Serine peptidases are involved in a number of essential intra- and extra-cellular functions, and have been directly implicated in pathogenic interactions between fungi and various hosts, including plants, with roles both in nutrient acquisition and immunity evasion [89, 90]. Sulfur metabolism is a core component of plant-pathogen interactions and plants secrete sulfur-rich molecules as a means of defense [91]. Vitamin B6/pyridoxine metabolism is linked to oxidative stress relevant for both plant defense and fungal pathogenicity [92, 93]. Serine hydrolase and antioxidant activities are among the significantly enriched functions of genes in the proximity of identified lncRNAs [Supplementary figure 4]. Regulation of hydrolase activity is the strongest enriched biological process among all potential targets compared to the rest of the genome [Supplementary figure 8], and the response to oxidative stress was the strongest enriched biological process for targets showing strong correlation with their interacting lncRNA [Supplementary figure 6].

**Figure 5:**
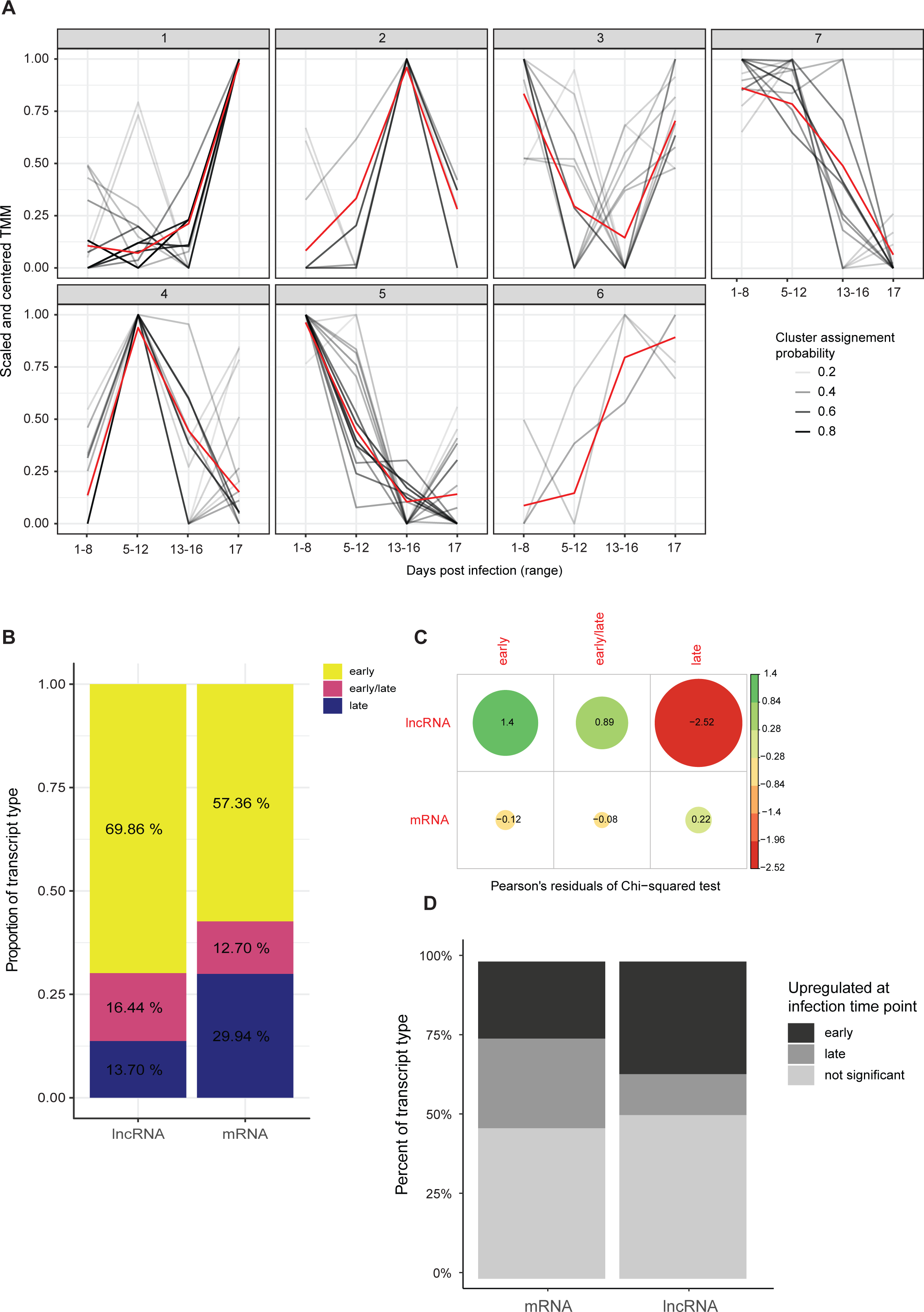
A) Trajectories of lncRNAs (black) across the infection cycle clustered using fuzzy c-means clustering. The y-axis represents the centered and scaled TMM values for each lncRNA. Lines in red show the cluster centers, which also take into account the trajectories of mRNAs. The grey-scale colored lines represent the confidence of assignment of each lncRNA into each cluster. Darker values show a higher confidence of belonging to the particular cluster. B) Distribution of lncRNAs and mRNAs into expression cluster types based on the peak expression of the cluster. Clusters in pink, defined as “early/late”, show bimodal peak expression, with high values during early and late infection, and lower values during the intermediate time points. C) Correlation plot showing the residuals for a Chi-squared test between the attribution of lncRNAs and mRNAs into cluster groups. The size of the dots represent the importance of the contributions of each condition and type. The color indicates the direction of the association (green for positive and red for negative). D) Percent of total lncRNAs and mRNAs differentially expressed either at to the early or late infection time point.

**Figure 6:**
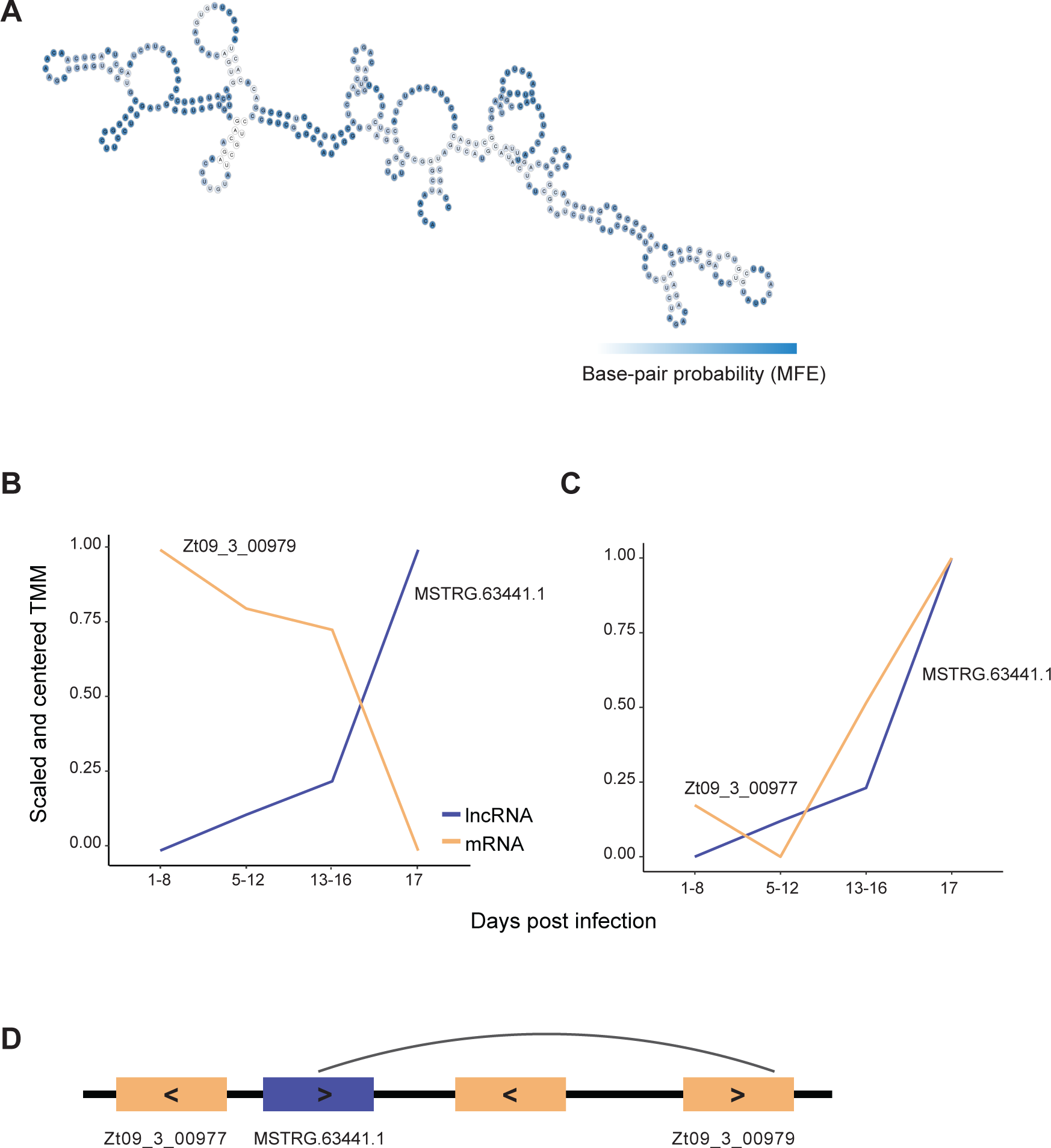
A) Secondary structure of the lncRNA MSTRG.63441.1 as predicted by free energy-minimization. The color shows the base pair probabilities. B) Trajectories across the infection cycle of the lncRNA MSTRG. 63441.1 (blue) and the nearby gene Zt09_3_00979 (orange). C) Ttrajectories across the infection cycle of the lncRNA MSTRG. 63441.1 and the nearby gene Zt09_3_00977. D) Schematic representation of the position of the lncRNA relative to the two genes. Arrows indicate the strand from which each element originates.

We assessed likely functionality of lncRNAs by testing for differential expression between early infection (time points A+B corresponding to 1-12 dpi) and late infection (C+D or 12-17 dpi). Transcripts were filtered for ≥10 reads across the 3 replicates retaining 62 lncRNAs. Nearly half (*n* = 30) out of which were found to be differentially expressed between early and late time-points (FDR 5%) [Supplementary table 3]. We found only 8 up-regulated lncRNAs compared to 22 lncRNAs down-regulated at the late infection stage [Figure 5D]. Of the 30 differentially expressed lncRNAs, 29 had an absolute correlation coefficient (Pearson’s *r*) greater than 0.8 with at least one gene within 50 kb [Supplementary table 4]. Moreover, 11 lncRNAs had at least one predicted interaction with a neighboring mRNA, of which 8 had a strong absolute (*r* > 0.8) correlation coefficient with their predicted mRNA target [Supplementary table 5].

We examined the differentially expressed lncRNAs and identified two showing strong expression correlations and predicted RNA interactions with nearby genes. MSTRG.9312.1 is upregulated during early infection and located ∼3 kb away from a predicted RiPP-like biosynthetic gene cluster (RiPP: ribosomally synthesized and post-translationally modified peptide) that contains 26 genes. MSTRG.9312.1 [Supplementary figure 13] is strongly correlated (*r* < -0.8; *r* >0.8) with 6 genes in this cluster [Supplementary table 6] and most significantly with *Zt09_7_00402* (*r* = 0.965; *p = 0.034*), from which the lncRNA is antisense downstream [Supplementary figure 14]. This gene encodes an uncharacterized protein with a DnaJ domain, typical of heat-shock proteins (HSPs), which are known to be implicated in various stress responses [94]. Additionally, MSTRG.9312.1 is predicted to be able to interact with the mRNA produced by the nearest same-sense neighbor, *Zt09_00404* (ndG = -0.1029) and is significantly positively correlated (*r*=0.961; *p* = 0.038).

The lncRNA MSTRG.6344.1 shows strong up-regulation during late infection (log fold-change = 2.08, *p* = 0.03) and a strong negative correlation with the expression of the gene *Zt09_03_00979* (*r* = -0.997; p = 0.0028) [Figure 7A-B]. *Zt09_03_00979* is one of the predicted mRNA targets of the lncRNA (ndG = - 0.101) and is located ∼5kb upstream on the same strand [Supplementary table 7]. *Zt09_3_00979* encodes an uncharacterized protein with an RNA recognition motif and a domain potentially interacting with the arginine methyl-transferase HMT1 (PRMT1 homolog), known to be implicated in chromatin dynamics through H4R3 methylation [95]. The activity of HMT1 on non-histone proteins has been shown to be important for virulence, growth and the response to stress in *F. graminearum* [96]. MSTRG.63441 is also strongly positively correlated, with *Zt09_03_00977* (*r* = 0.91) [Figure 7C] which encodes an uncharacterized protein with an OTT_1508-like deaminase domain. The gene is located ∼700 bp upstream of the lncRNA transcribed from the opposite strand and in the opposite direction. The the nearest upstream gene to the lncRNA is *Zt09_03_00978* with which it shows no transcriptional correlation [Figure 7D].

**Figure 7:**
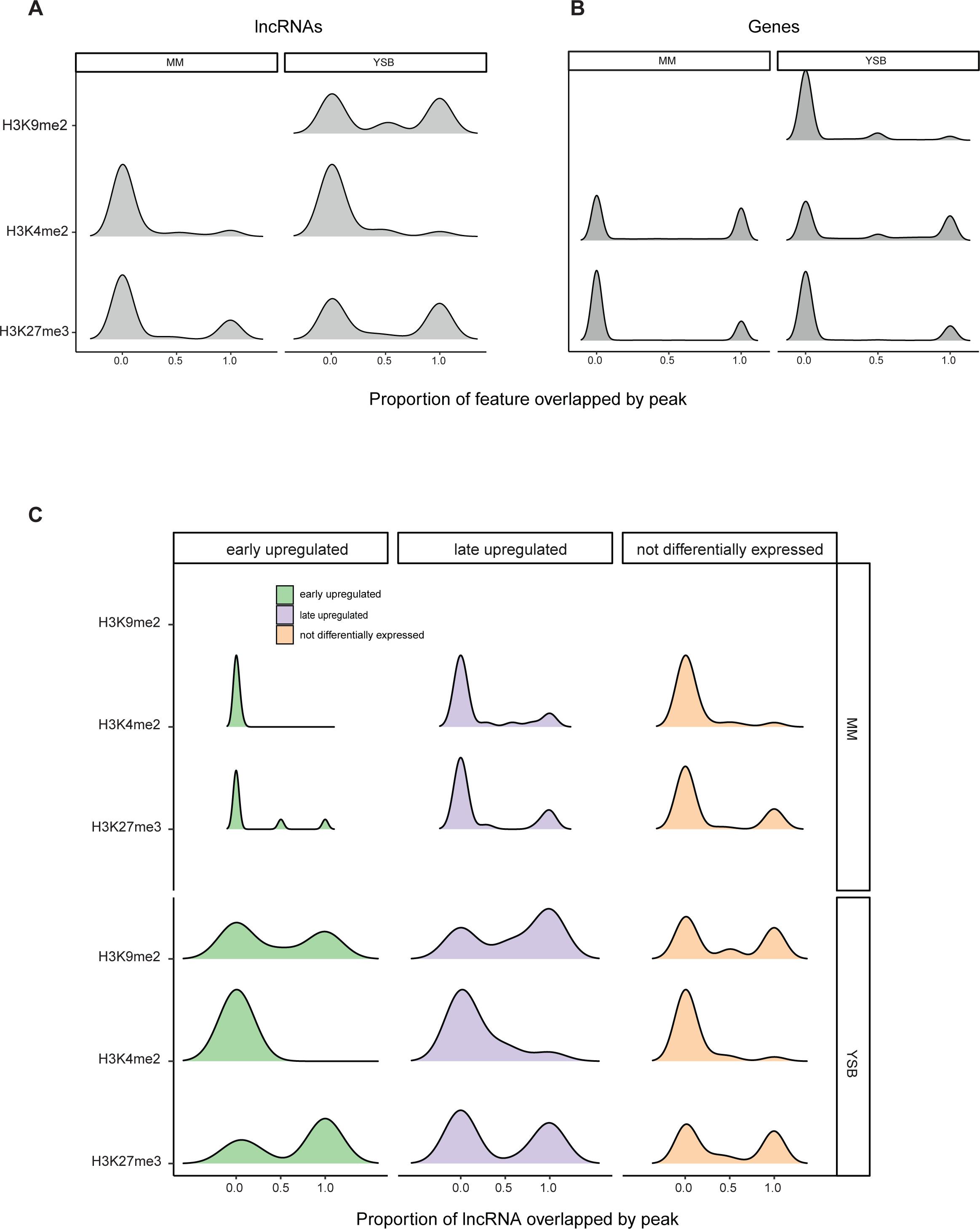
Density plot showing the coverage of (A) lncRNAs and (B) genes according to different histone methylation markers in minimal-media (MM) and yeast-sucrose broth (YSB). MM mimics conditions during early infection. The proportion of each locus covered by the marker is shown. C) Density plot comparing the coverage of lncRNAs by different histone methylation marks based on the differential expression status of the lncRNAs and the time point of their peak expression in MM and in YSB.

### Epigenetic states of lncRNA loci

In *Z. tritici,* genes encoding virulence factors are most likely regulated through histone modification [97]. Notably, heterochromatin associated with H3K27me3 and H3K9me3 is enriched in both species-specific and biosynthetic genes, whereas euchromatin associated with H3K4me2 covers mostly gene-dense and conserved regions [97]. Data for H3K27me3, H3K9me3, and H3K4me2 [98] for the reference isolate IPO323 grown in both minimal media (MM), and H3K4me2 and H3K27me3 data for growth in YSB was analyzed. Compared to genes, lncRNAs were less likely to be covered by H3K4me2 in both growth conditions (Welch two-sided *t*-tests; *p*-values < 2.2e-16) [Figure 7A-B]. LncRNAs were more likely to be covered by H3K9me2 in YSB (Welch two-sided *t-*test, *p*-value = 1.007e-10), and H3K27me3 in both MM and YSB (Welch two-sided *t-*test, *p*-value = 2.585e-07) compared to genes [Figure 7A-B]. Interestingly, lncRNA loci but not genes showed an increase in H3K27m3 coverage in YSB compared to MM [Figure 7A]. The higher proportion of lncRNAs covered by H3K27me3 compared to genes is consistent with the notion that lncRNAs are likely to be species-specific.

Soyer et al. [97] showed that *in vitro* H3K27me3 and H3K9me3 marked regions were enriched in genes that were differentially expressed during early host colonization or at the switch to necrotrophic growth. We compared *in vitro* chromatin profiles of lncRNAs upregulated during either early infection or late infection. In MM, we find that lncRNAs upregulated during early infection lack coverage by H3K4me2, and are rarely covered by H3K27me3 (1 out of 22 early upregulated lncRNAs are fully covered by H3K27me3 in MM). In contrast, lncRNAs upregulated during late infection show most coverage by H3K7me3 (4 out of 8 late upregulated lncRNAs are fully covered by H3K27me3 in MM) [Figure 7C]. In YSB, all lncRNAs show an increase in coverage by H3K27me3. The MM conditions *in vitro* resemble conditions during early growth *in planta,* due to the lack of external resources and higher levels of stress [49]. Hence, the lncRNAs involved in early infection may be epigenetically regulated and repressed by H3K27m3.

### Pangenome analyses of lncRNA diversity

*Z. tritici* populations are highly diverse even at small geographic scales [42, 99, 100], and the species carries a vast accessory genome [61, 101, 102]. We aimed to understand if lncRNAs were as diverse as genes across the pangenome of the species. In order to compare gene diversity to lncRNA diversity, we repeated our prediction pipeline using transcriptomic data from four additional isolates (1A5, 1E4, 3D1 and 3D7), collected at similar time points during the infection cycle as the predictions based on Zt09. We clustered all predicted lncRNA transcripts using low stringency (sequence identity ≥80%, minimum alignment coverage 75%) to account for the fact that lncRNA sequences may have diverged more rapidly than protein-coding sequences [103]. We obtained 1671 clusters of lncRNAs, of which 1233 contained only a single transcript. We constructed an accumulation curve using the cluster attributions [Figure 8A] and compared this to the accumulation of gene orthogroups as previously constructed for the same isolates [101] [Figure 8B]. Compared to genes, the slope of the lncRNA accumulation curve was steep and without any indication of a plateau, suggesting that lncRNAs are more diverse than genes. However, many lncRNAs show highly specific expression patterns and may only be expressed during particular stages of development and conditions [103]. Condition specificity is much higher than for genes [26]. Considering that the data for 1A5, 1E4, 3D1 and 3D7 were taken at the same time-points regardless of morphological characteristics, some of the observed lncRNA diversity may be due to small variation in developmental stages at identical time-points.

**Figure 8:**
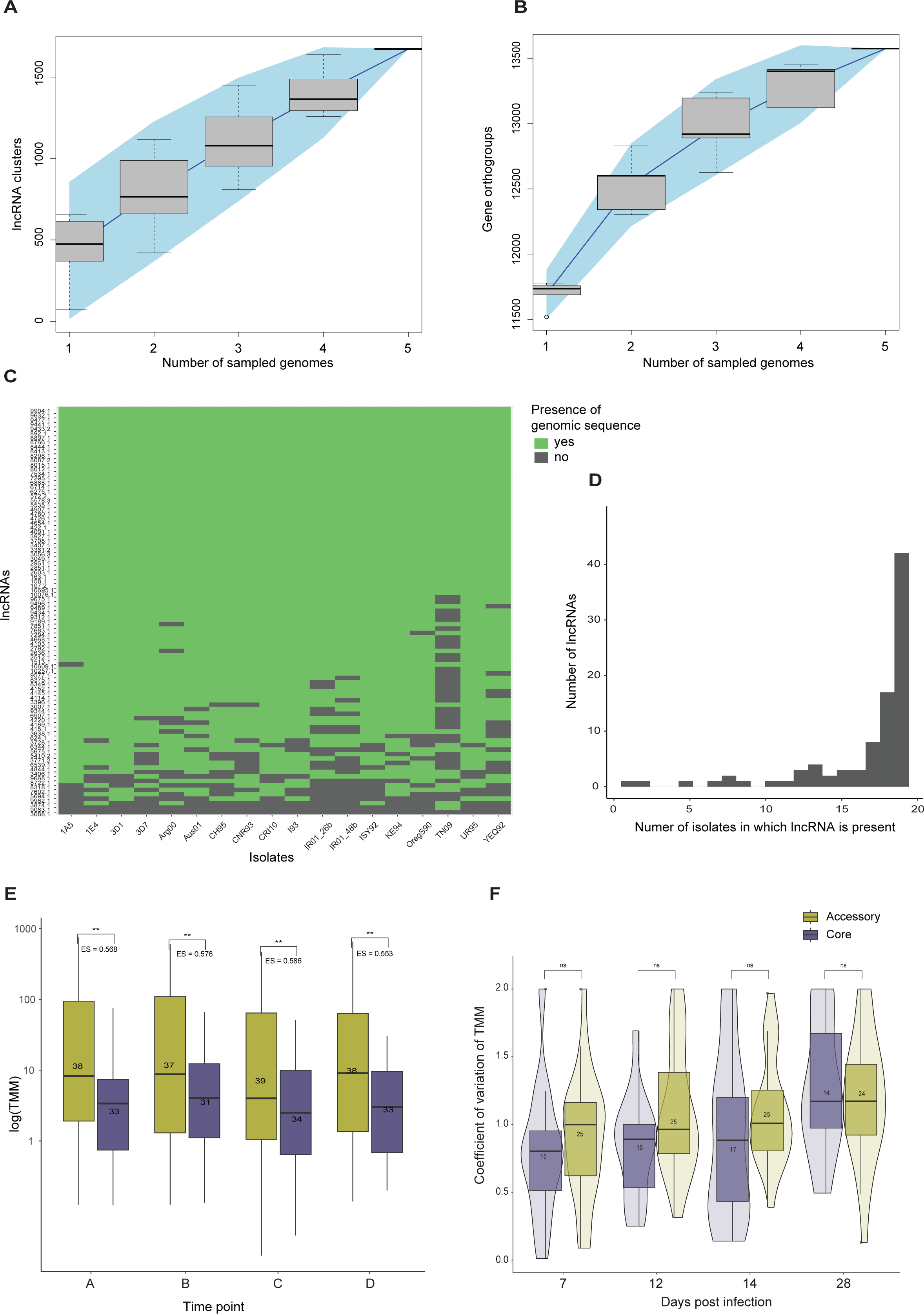
A) Accumulation curve of lncRNAs predicted using transcriptomic data for four additional reference strains clustered based on sequence similarity. Genomes were sampled without replacement. B) Accumulation curve of gene orthogroups. Genomes were sampled without replacement. C) Heatmap showing the of presence of each lncRNA predicted in 19 different isolates at the genomic level. If genomic sequences are identified to be similar to the lncRNA reported in Zt09, the heatmap is colored in green. D) Bar plot showing the number of isolates in which lncRNAs of Zt09 were found with a match at the DNA level. E) Comparison of expression values (TMM) between core lncRNAs (found in all genomes) and accessory lncRNAs (missing in at least one genome), for all lncRNAs in the strain Zt09 at different time-points during the infection cycle. Stars represent the significance level of Welch two-sided *t-*tests comparing mean expression between core and accessory lncRNAs. Values denoted by “ES=” show the effect size relative to the expression of accessory lncRNAs. Numbers in each box show the sample sizes. F) Violin plot comparing the coefficient of variation (CV) in expression between core and accessory lncRNAs, calculated by comparing centered and scaled TMM values between four isolates across the infection cycle. Numbers inside the boxes show the sample size in each group.

To assess whether the observed lncRNA diversity may be linked to specific conditions or expression variability, we performed a sequence similarity based search for each lncRNA identified in Zt09 in the 18 additional reference quality genome assemblies including the four used in the previous section [Figure 8C]. In contrast to the transcriptome-based approach, we found that the overwhelming majority of lncRNA sequences were conserved among all 19 genomes [Figure 8D] and only a single lncRNA was found uniquely in Zt09. Hence, the pool of expressed lncRNAs likely varies greatly between strains even at similar time points. We used genetic similarity to assign core (present in at least 18 out of 19 isolates) and accessory status for each lncRNA identified in Zt09. Core lncRNAs are expressed at significantly lower levels than accessory lncRNAs at all time points [average effect size of 0.570] [Figure 8E]. Significant differences were also found when comparing expression levels of core and accessory lncRNAs in the strains 1A5, 1E4, 3D1, and 3D7, except at 7 days post infection [Supplementary figure 15]. Low levels of lncRNA expression may be an important feature of lncRNA functionality, notably for those involved in chromatin organization [103, 104]. Compared to accessory lncRNAs, core lncRNAs tends to have smaller coefficients of variation in their expression levels between isolates, although these differences were not significant [Figure 8F]. Furthermore, core and accessory lncRNAs show no distinct distribution (Chisq-test *p*-value = 0.24) among expression clusters [Supplementary figure 16], despite the fact that accessory lncRNAs tend to be underrepresented in clusters with peak expression at later time-points [Supplementary figure 17].

## Discussion

We identified lncRNAs in the fungal pathogen *Z. tritici*, and evaluated potential regulatory and biological functions. Compared to similar genome-wide screens for lncRNA candidates in other organisms, including studies on other pathogenic fungi, our prediction yielded very low numbers of lncRNA candidates. Most studies report 1000-10,000s lncRNAs per species [20, 34, 35, 38, 105]. We hypothesize that this results from a high number of false negatives. We chose to use a higher weight on the Youden’s index than the default parameter, which is used in the prediction tool to determine a cut-off for sensitivity. A higher weight improves false-positive rates but increases false-negatives[54]. Additionally, only a single transcript per locus was considered, despite the fact that lncRNAs can undergo alternative splicing [19, 106, 107]. The choice to include TE consensus sequences in the known set of mRNAs may have also increased false negatives considering that many lncRNAs may contain TE-associated elements [108]. Lastly, some lncRNAs may have been missed due to the single-ended nature of our RNA-seq data, considering that paired-end reads increase library complexity and result in higher read counts per locus, particularly in non-coding regions, which improves transcript assembly [109]. Because we focused on *in planta* expression, a low proportion of the sequenced biological material originates from the pathogen, which results in low read counts. In such cases, the increased library complexity offered by paired-end sequencing may be of particular importance. Paired-end data can also improve the detection of overlapping same-sense transcripts [110], which may be the case for a substantial fraction of fungal lncRNAs. In *M. robertsii,* 12% of identified lncRNAs share exons with known mRNAs [34].

Although many fungal lncRNAs are intergenic [34, 35, 38], evidence from several fungal pathogens suggests that a large proportion of their lncRNAs could be antisense to protein-coding genes [35, 37, 111]. A genome-wide study in the fungal pathogen *C. neoformans* showed that >60% of predicted lncRNAs were antisense [37], in *M. robertsii* 49% [111], and in *U. virens* 33% [35]. Surprisingly, only two antisense lncRNAs were identified using our approach. Comparison with the new annotation of the reference genome [52], in which similar numbers of sense and antisense lncRNAs were found, suggests that technical issues are the most likely cause of the lack of antisense lncRNA in our predictions. In some cases, reads originating from the loci in the new annotation are present but at low depth, and/or they do not cover the full length of the transcript [52]. As such, transcripts were not properly assembled and were never assessed by the prediction tool. In other cases, transcripts were assembled correctly but were predicted as coding, perhaps because of shared features with the opposing coding sequence, which may make antisense lncRNA more challenging to distinguish from mRNA compared to intergenic lncRNA.

LncRNAs showed differences in epigenetic profiles compared to genes, and were more likely to be located in facultative heterochromatic regions. In *Z. tritici*, these regions are associated with effectors and biosynthetic genes, which are often up-regulated during early colonization [97]. The presence of lncRNAs in facultative heterochromatic regions may be indicative of their co-regulation with genes found in the same regions, which is consistent with the rest of the observations in this study. The fact that lncRNAs with peak expression during late infection were more likely to be covered by repressive markers in minimal media than those with peak expression at earlier times points, suggests that lncRNA expression and chromatin state are linked. LncRNAs are known to contribute to the formation of heterochromatin in *Drosophila* and plants through interactions with chromatin modifying enzymes [112]. A candidate lncRNA shows the potential to interact with HMT1, through a domain in a neighboring, anti-correlated and potentially chemically targeted mRNA. The interaction is unlikely to be directly related to the formation of heterochromatin itself, however it demonstrates the potential for lncRNAs to be involved in chromatin dynamics in this species. As little is known about the mechanisms of epigenetic regulation in *Z. tritici,* it may be interesting for future research to consider the role of lncRNAs, especially in dynamically regulated regions.

LncRNAs were expressed at lower levels than mRNAs, which is consistent lncRNA expression in other organisms. As expected, expression is also generally more variable between isolates, except at 7 days post infection. It is important to note that this estimate is susceptible to overestimation due to low read counts and high stochasticity. We find no evidence that the lncRNA position relative to a gene influences the level of co-expression between the pair, showing that lncRNAs are neither predominantly *cis*- or *trans*-acting in this species. The potential for lncRNAs to form RNA-complexes with nearby mRNAs does significantly impact anti-correlation between lncRNA and mRNA expression at close distances. This suggests that lncRNA-mRNA interactions in *cis* may be one mechanism of lncRNA-mediated regulation. The majority of identified lncRNAs were upregulated during early stages of host infection. Compared to genes, a significantly larger proportion of lncRNAs show expression profiles with peaks either before or during the switch to necrotrophic growth. Moreover, their location in specific chromatin regions is consistent with that of protein coding genes expressed at these time-points. Interestingly, this mirrors observations from the human microbial parasite *Cryptosporidium parvum* with lncRNAs being highly expressed during invasion and less during proliferation in contrast to mRNAs [33]. During early colonization, the pathogen must rely on internal nutrient stores, as the organism is unable to acquire these directly from the host [85]. The pathogen must also protect itself from host-defense mechanisms, such as the production of reactive oxygen species and chitinases [89, 113]. LncRNAs are known to regulate a wide array of stress-response pathways and changes in nutrient acquisition in other organisms [21, 32]. Hence, lncRNAs may be of particular importance during this life cycle stage. The idea that lncRNAs are involved in responding to early-infection stress in *Z. tritici* is supported by enriched functions among potential lncRNA targets, such as antioxidant or serine-hydrolase activity.

*Z. tritici* is a highly polymorphic species including at the level of gene presence-absence variation and accessory chromosomes [101]. We explored pangenomic variation of lncRNAs in comparison to genes. Most lncRNAs were located on the core chromosomes shared between all individuals of the species. Nevertheless, based on the transcriptomes of four reference isolates, we observe a high level of variation in lncRNA repertoires, and a steep accumulation curve compared to genes. At the genomic level, the majority of lncRNAs identified in the reference strain IPO323 are shared among 18 strains from around the world. Differences in the genomic and transcriptomic evidence of lncRNAs are best explained by the high specificity of lncRNAs [26]. LncRNA expression is highly dependent on environmental conditions, and small changes in developmental stage of either the pathogen or the host, or in experimental conditions, may result in vastly different lncRNA repertoires being expressed. While similar loci are shared among isolates, mutations may have accumulated in these loci to affect how lncRNA are transcribed. Future studies should aim to disentangle transcriptomic diversity from genomic diversity, particularly in the context of lncRNAs. In conclusion, our study provides a repertoire of lncRNAs loci for *Z. tritici* and how these likely intervene in host infection processes and the responses to stress.

## Acknowledgements

We thank Nicolas Lapalu and Marc-Henri Lebrun for providing early access to a long-read transcriptomic dataset for IPO323.

