## Supplementary Figures for "The expression landscape and pangenome of long non-coding RNA in the fungal wheat pathogen *Zymoseptoria tritici*"

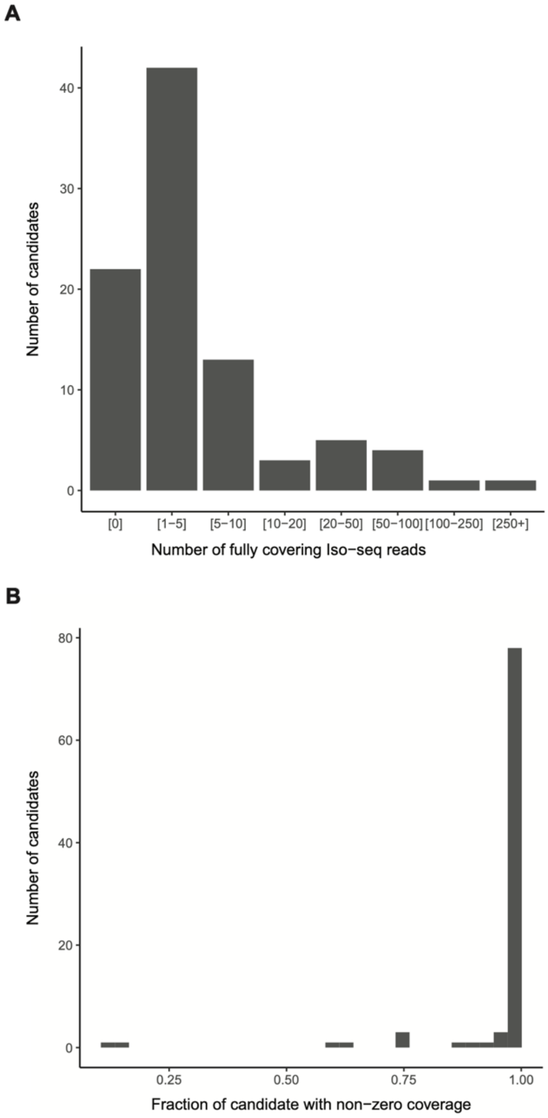


**Supplementary figure 1:** A) Bar chart summarizing the number of fully covering unique Iso-seq reads per lncRNA candidate predicted in the current study. The y-axis shows the number of lncRNA candidates that have the number of fully covering Iso-seq reads represented in each bin on the x-axis. B) Histogram showing the distribution of coverage of lncRNA candidates by pooled Iso-seq reads. The x-axis represents the proportion in length of each candidate that is covered by pooled iso-seq reads. A fraction of 1 means that the entire length of the candidate is covered, although reads may be non-continuous.

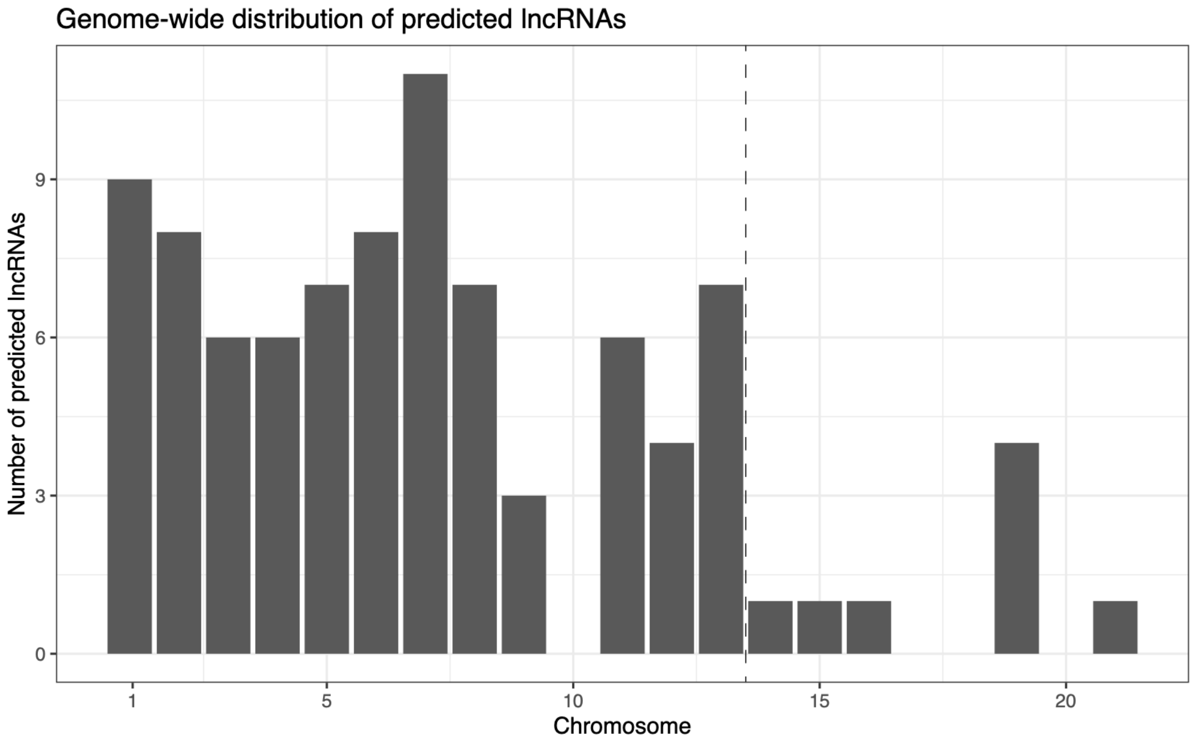


**Supplementary figure 2:** Distribution of the number of lncRNA candidates on each chromosome. The vertical line shows the separation between core chromosomes, on the left, and accessory chromosomes on the right. The y-axis shows the count of lncRNA candidates on each chromosome and is not normalized by chromosome length.

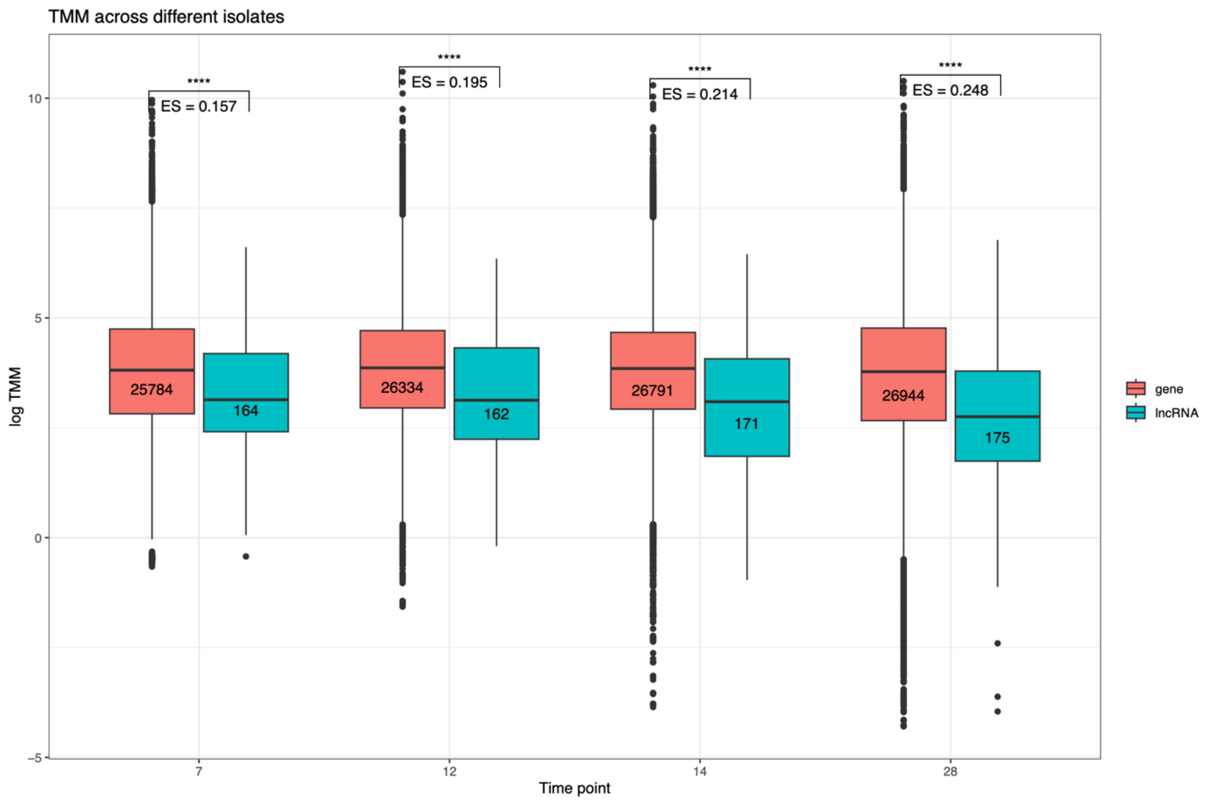


**Supplementary figure 3:** Box-plots comparing expression in log transcripts-per-million (TMM) between lncRNAs (blue) and mRNAs (red) for 4 additional reference strains. Numbers inside the boxes show the sample size for each group. Stars represent the significance level concerning the difference between expression of lncRNAs and mRNAs at each time point (indicated by parentheses), tested using a two-sided Welch *t*-test. Each time-point and each isolate was tested individually.

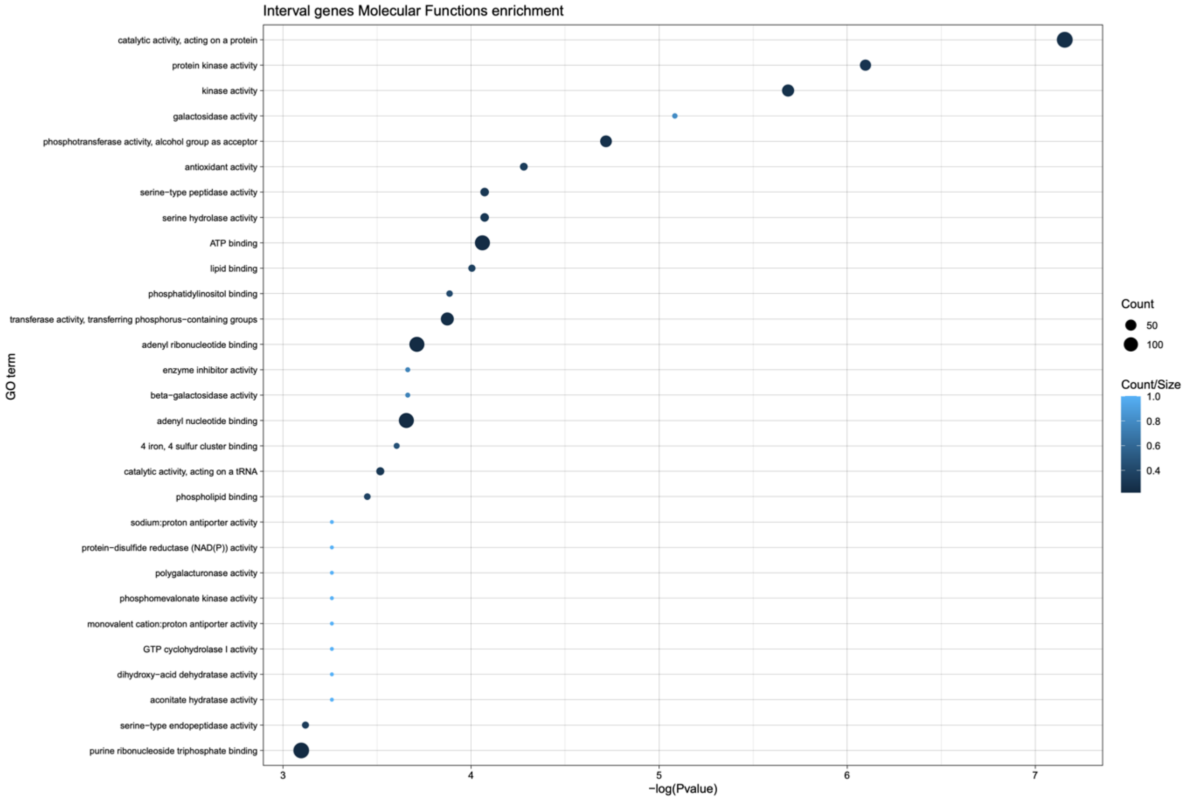


**Supplementary figure 4:** Plot summarizing the significant results of an enrichment test against the genome for molecular function, for genes located within 50 kb of a lncRNA candidate. The y-axis describes the enriched functions, and the x-axis shows the -log p-value of the enrichment. A higher value on the x-axis means higher statistical significance. The size of each dot represents the number of genes belonging to the group of interest in each functional category. A larger dot indicates that multiple genes with a particular function are found within 50 kb of a lncRNA. The color of the the dot represents the number of genes belonging to the group of interest in each category, divided by the genome-wide total. A light dot means that the majority of all genes annotated with a particular function are found within 50 kb of a lncRNA.

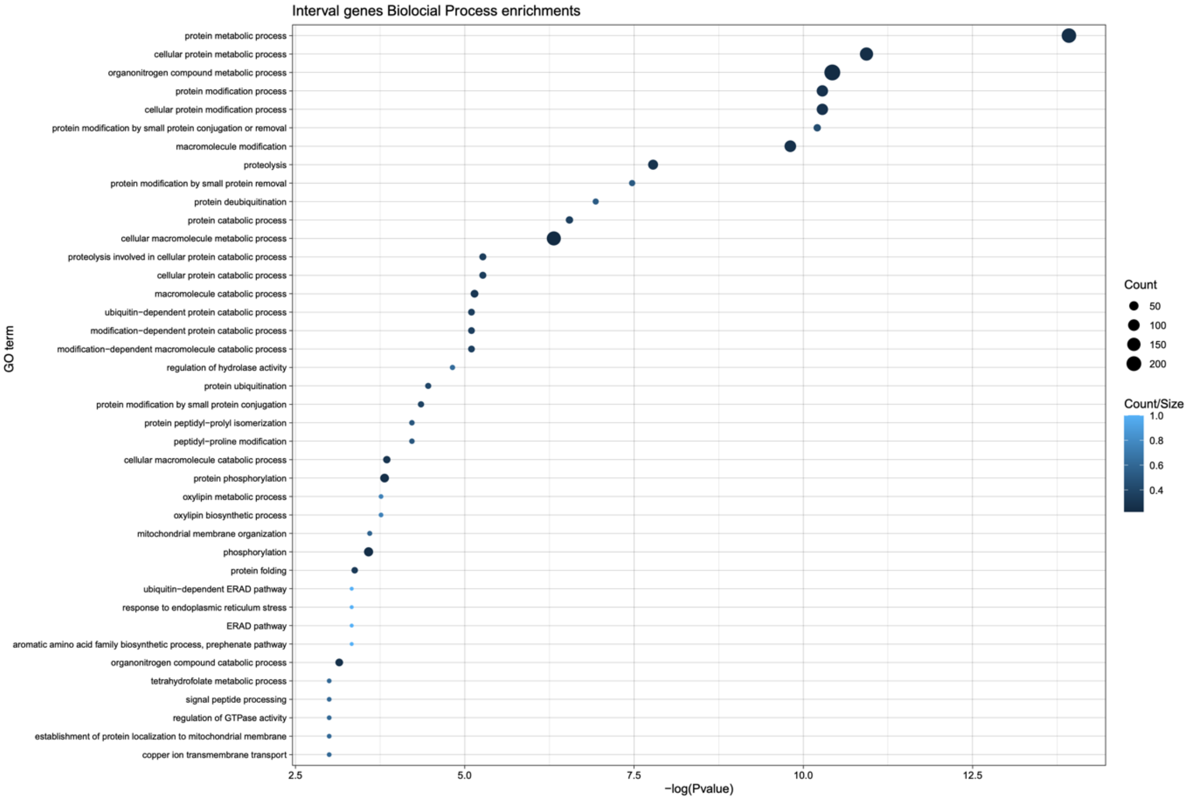


**Supplementary figure 5:** Plot summarizing the significant results of an enrichment test against the genome for biological processes, for genes located within 50 kb of a lncRNA candidate. The y-axis describes the enriched functions, and the x-axis shows the -log p-value of the enrichment. A higher value on the x-axis means higher statistical significance. The size of each dot represents the number of genes belonging to the group of interest in each functional category. A larger dot indicates that multiple genes with a particular function are found within 50 kb of a lncRNA. The color of the the dot represents the number of genes belonging to the group of interest in each category, divided by the genome-wide total. A light dot means that the majority of all genes annotated with a particular function are found within 50 kb of a lncRNA.

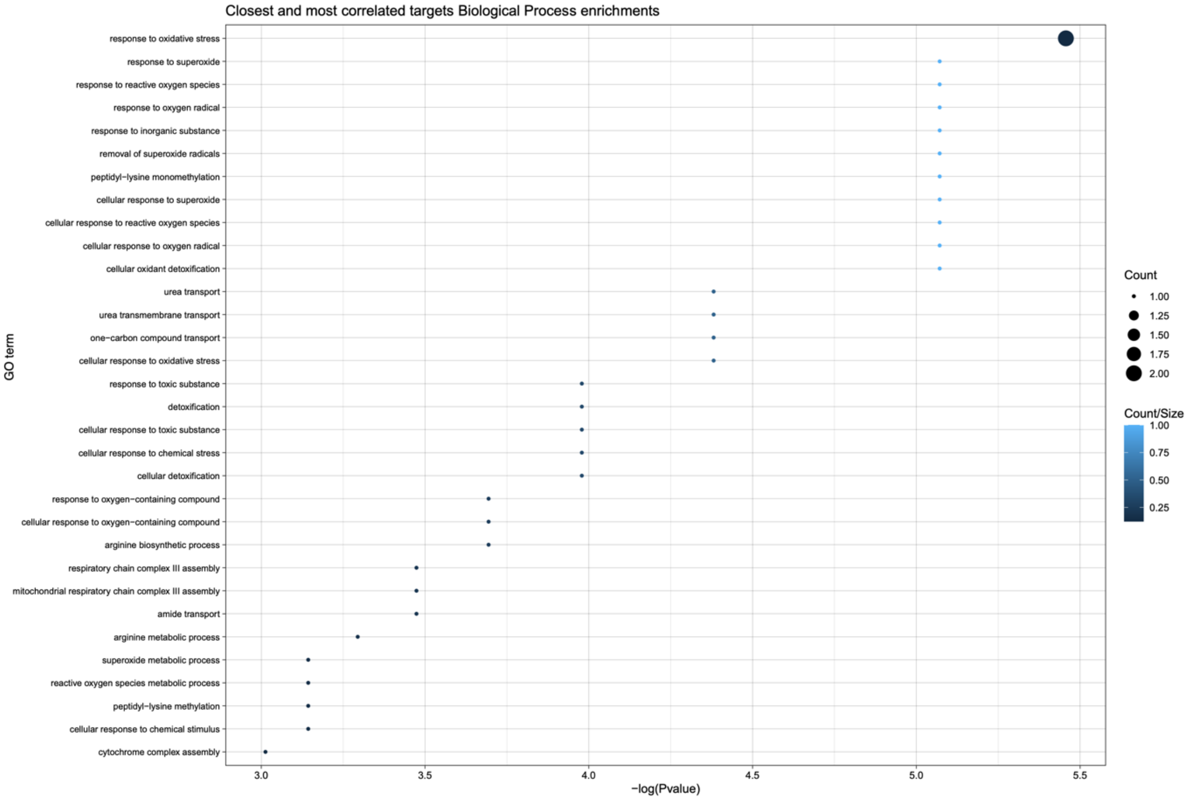


**Supplementary figure 6:** Plot summarizing the significant results of an enrichment test against the genome for biological processes, for genes located within 50 kb of a lncRNA candidate that also show strong expression correlation (absolute Pearson's $r > 0.8$) with said candidate. The y-axis describes the enriched functions, and the x-axis shows the -log p-value of the enrichment. A higher value on the x-axis means higher statistical significance. The size of each dot represents the number of genes belonging to the group of interest in each functional category. The color of the the dot represents the number of genes belonging to the group of interest in each category, divided by the genome-wide total.

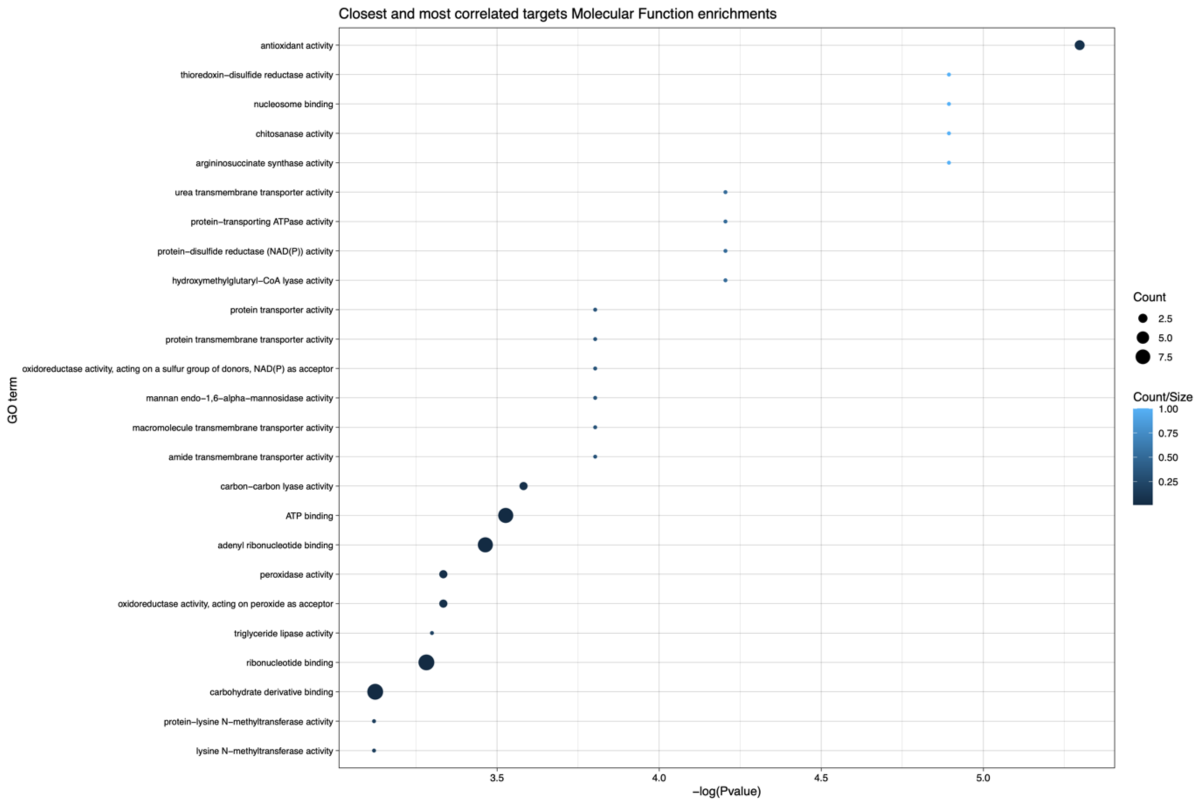


**Supplementary figure 7:** Plot summarizing the significant results of an enrichment test for molecular function, for genes located within 50 kb of a lncRNA candidate that also show strong expression correlation (absolute Pearson's $r > 0.8$) with said candidate. The y-axis describes the enriched functions, and the x-axis shows the -log p-value of the enrichment. A higher value on the x-axis means higher statistical significance. The size of each dot represents the number of genes belonging to the group of interest in each functional category. The color of the the dot represents the number of genes belonging to the group of interest in each category, divided by the genome-wide total.

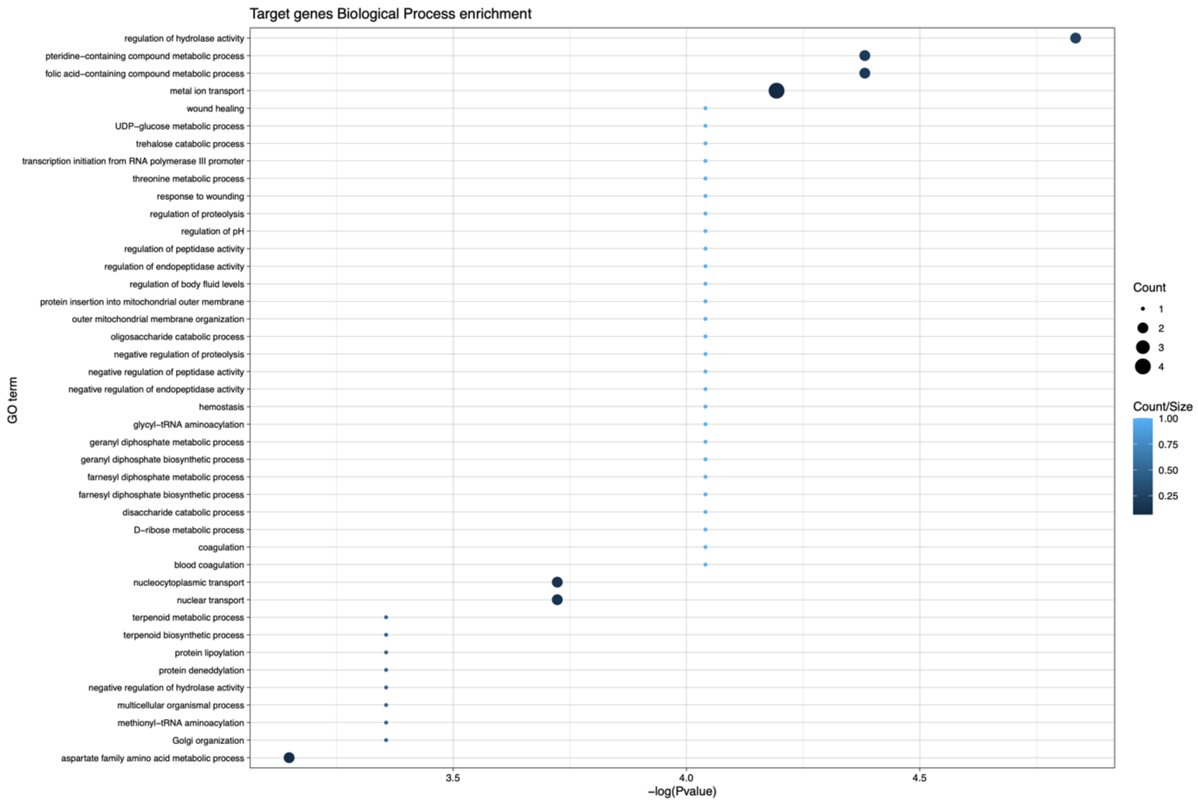


**Supplementary figure 8:** Plot summarizing the significant results of an enrichment test against the genome for biological processes of potential *cis*-RNA-targets of lncRNA candidates. The y-axis describes the enriched functions, and the x-axis shows the -log p-value of the enrichment. A higher value on the x-axis means higher statistical significance. The size of each dot represents the number of genes belonging to the group of interest in each functional category. The color of the the dot represents the number of genes belonging to the group of interest in each category, divided by the genome-wide total.

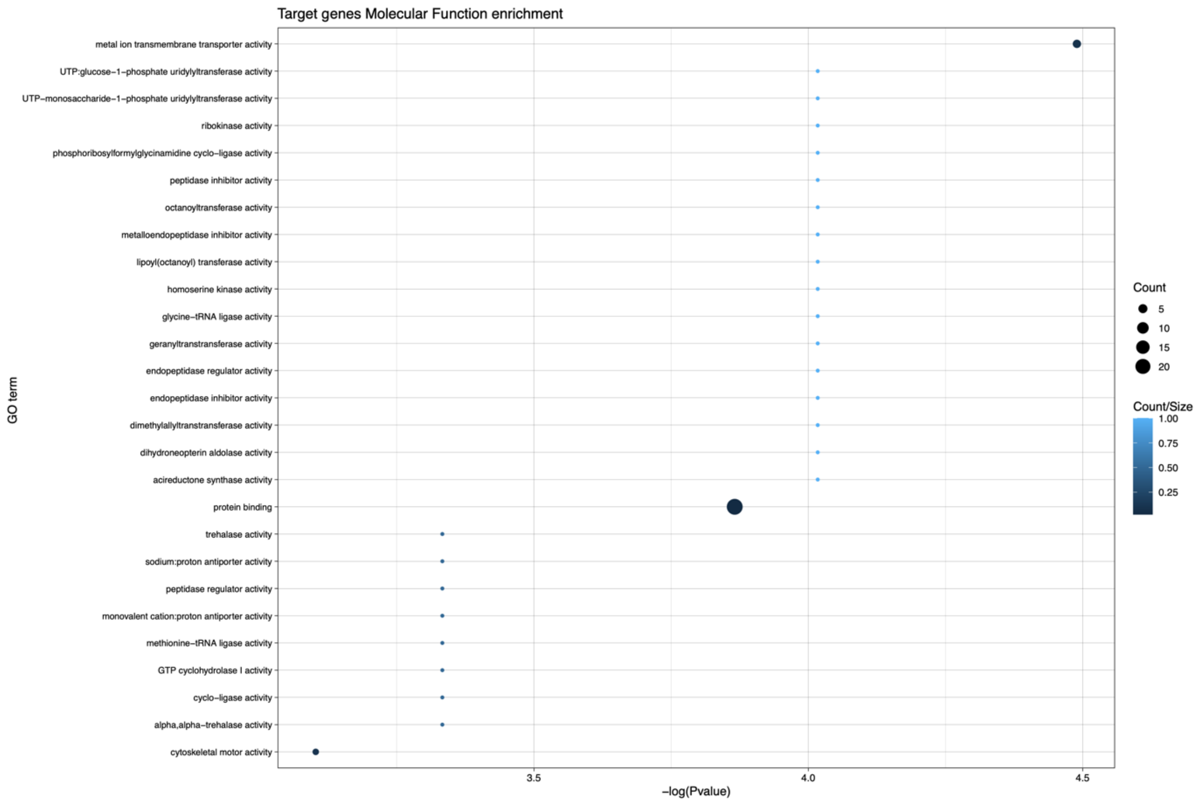


**Supplementary figure 9:** Plot summarizing the significant results of an enrichment test against the genome for molecular function of potential *cis*-RNA-targets of lncRNA candidates. The y-axis describes the enriched functions, and the x-axis shows the -log p-value of the enrichment. A higher value on the x-axis means higher statistical significance. The size of each dot represents the number of genes belonging to the group of interest in each functional category. The color of the the dot represents the number of genes belonging to the group of interest in each category, divided by the genome-wide total.

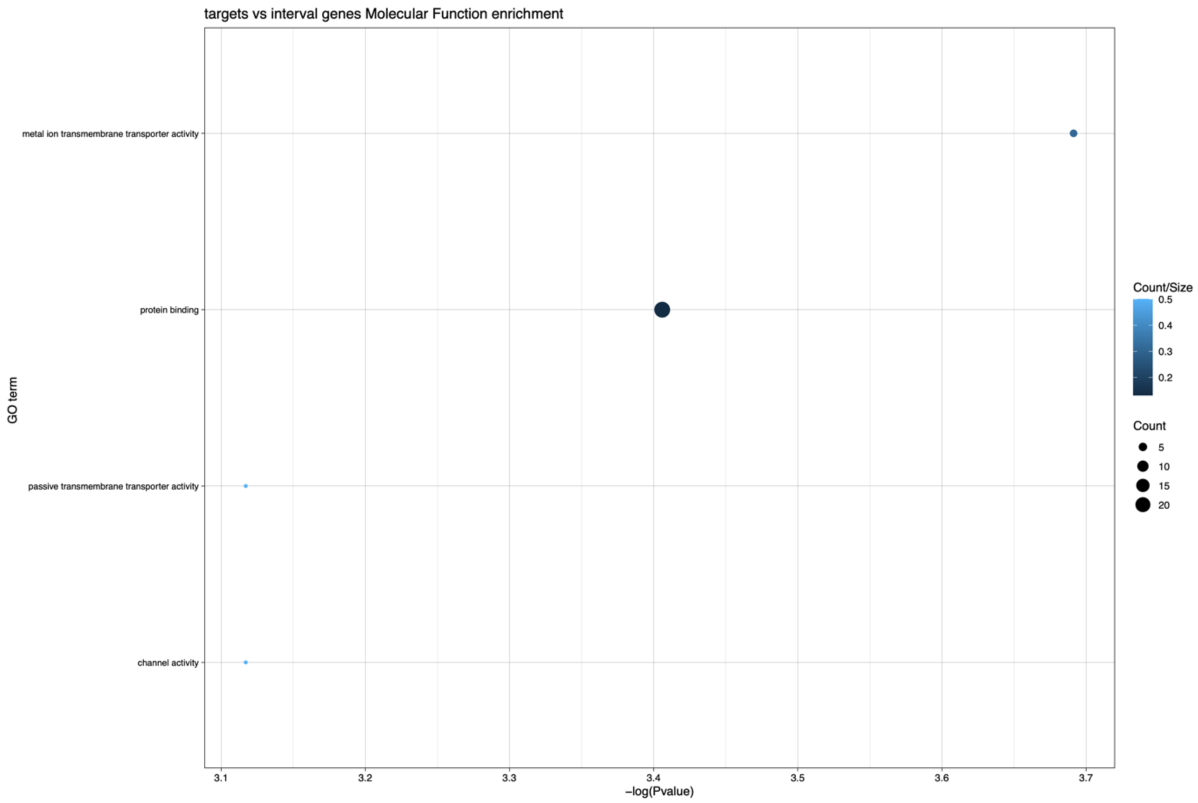


**Supplementary figure 10:** Plot summarizing the significant results of an enrichment test against all genes within 50 kb of a lncRNA candidate, for molecular function, of potential *cis*-RNA-targets of lncRNA candidates. The size of each dot represents the number of genes belonging to the group of interest in each functional category. The color of the the dot represents the number of genes belonging to the group of interest in each category, divided by the total number of genes in each category within a 50 kb interval around a lncRNA candidate.

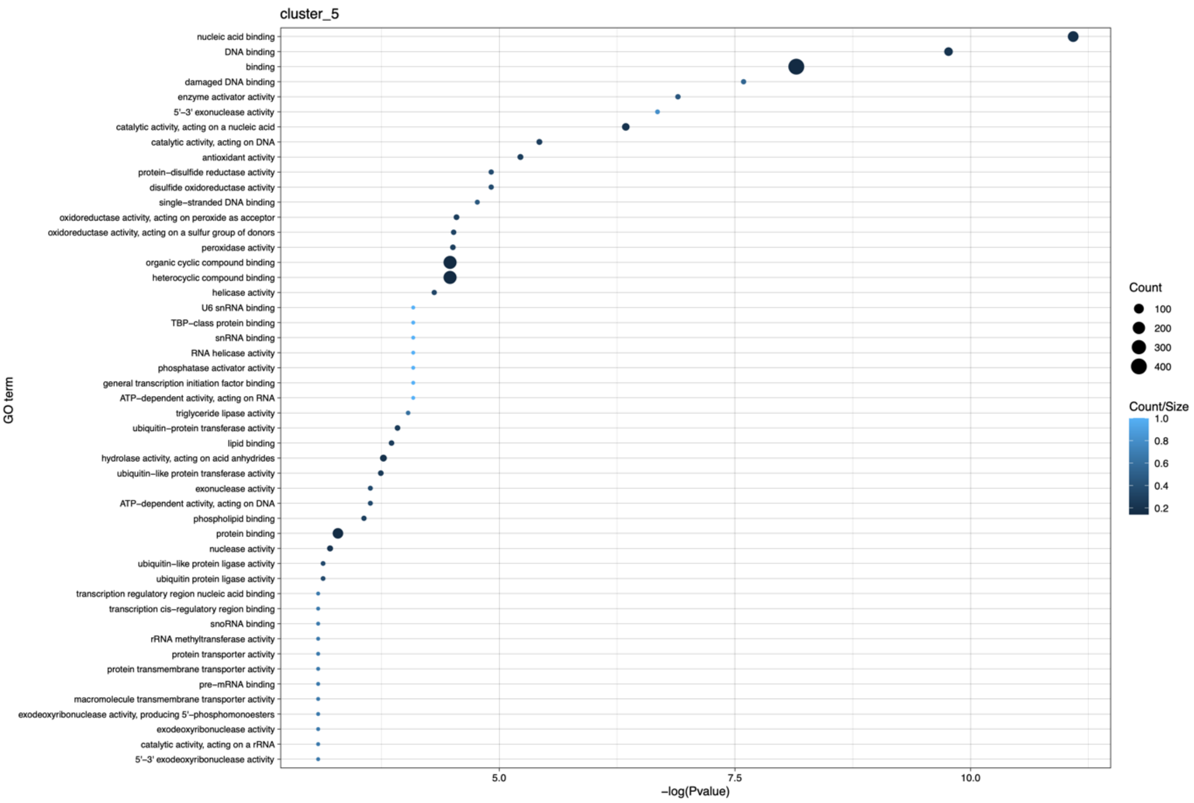


**Supplementary figure 11:** Plot summarizing the significant results of an enrichment test against the genome for molecular function of all genes belonging to the expression cluster 5 (which shows peak expression a 1 day post infection (dpi) and contains the highest number of lncRNAs). The size of each dot represents the number of genes belonging to the group of interest in each functional category. The color of the the dot represents the number of genes belonging to the group of interest in each category, divided by the genome-wide total.

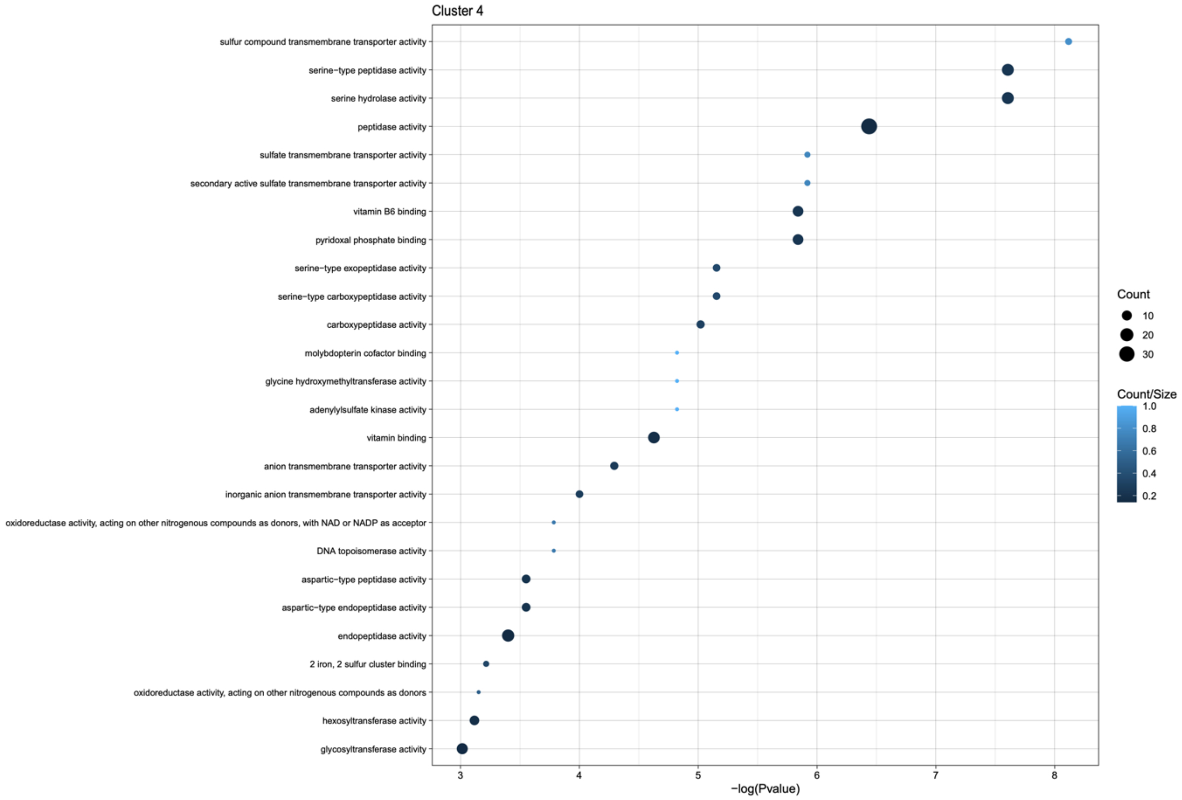


**Supplementary figure 12:** Plot summarizing the significant results of an enrichment test against the genome for molecular function of all genes belonging to the expression cluster 4 (which shows peak expression at 5-12 day post infection (dpi), and the second highest number of lncRNAs). The size of each dot represents the number of genes belonging to the group of interest in each functional category. The color of the the dot represents the number of genes belonging to the group of interest in each category, divided by the genome-wide total.

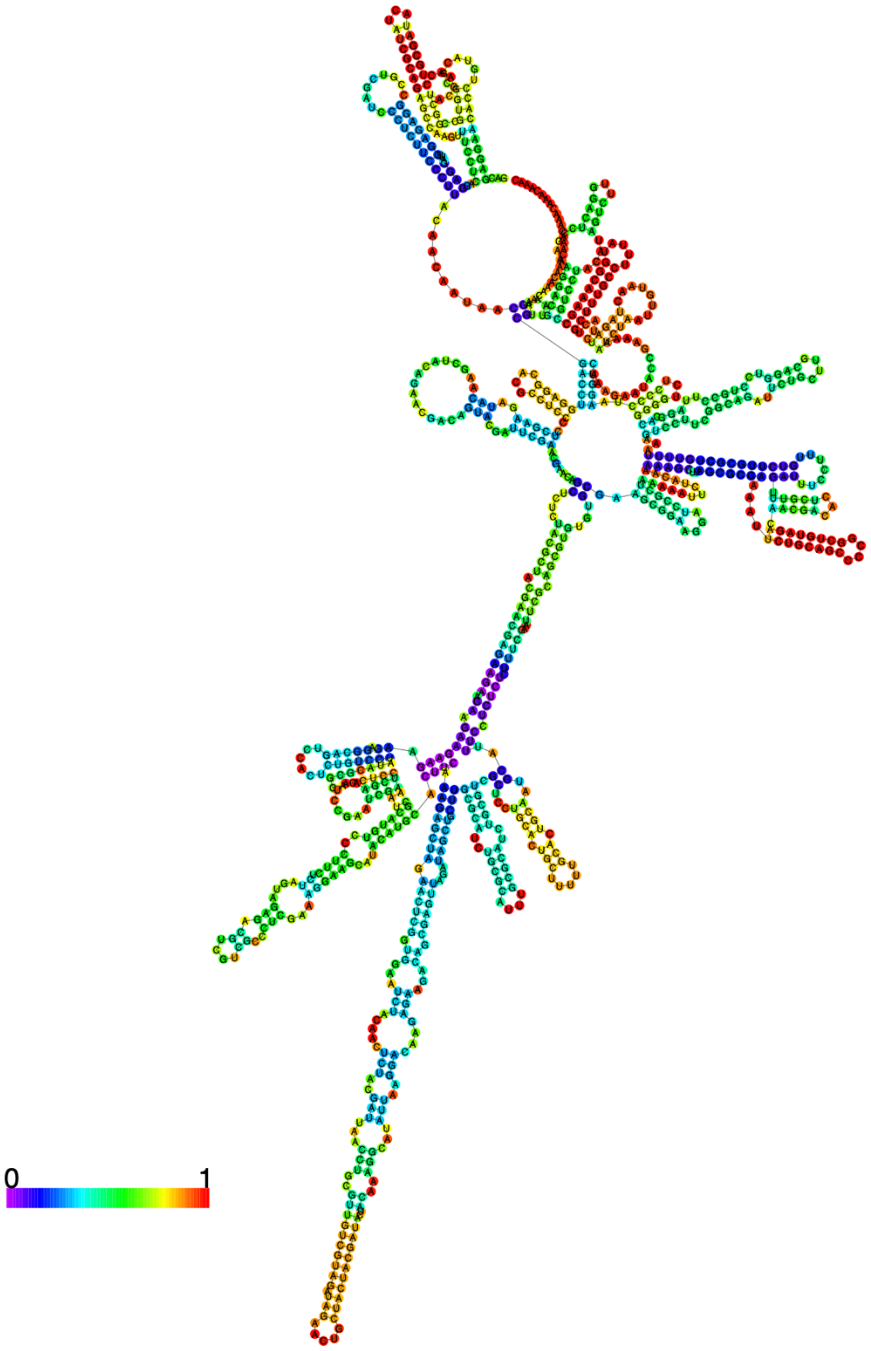


**Supplementary figure 13:** Predicted secondary structure of the lncRNA candidate MSTRG.9312.1, as output by RNA-fold using MFE. The color of the dots represents the base-pair probabilities at each position, with warmer colors indicating a higher probability.

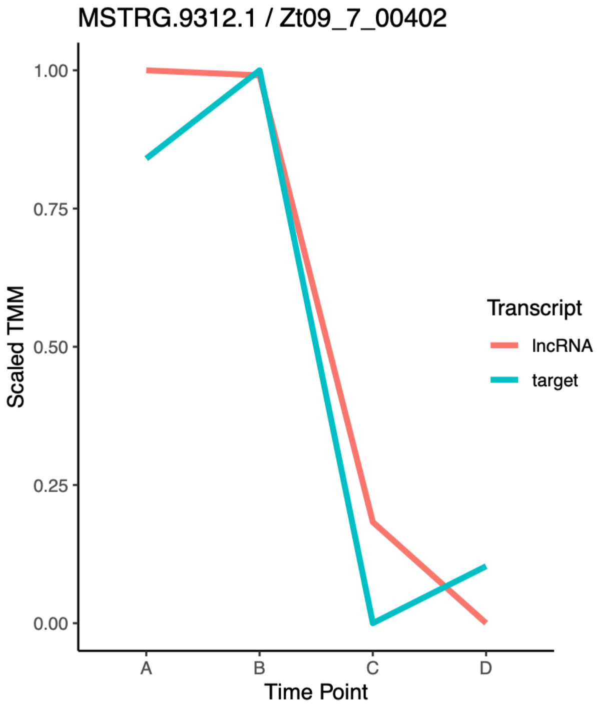


**Supplementary figure 14:** Expression trajectories across the infection cycle of a gene (*Zt09_7_00402*) encoding a predicated DnaJ domain (blue), and the downstream antisense lncRNA MSTRG.9312.1 (red). Both loci are located within a predicted RiPP cluster. The y-axis shows the scaled and centered TMM values for better comparability between the two transcripts.

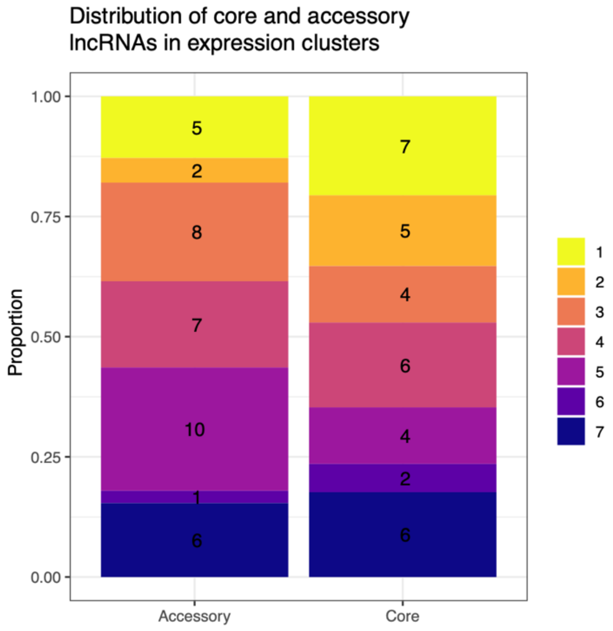


**Supplementary figure 15:** Bar chart summarizing the distribution of lncRNA candidates into each expression cluster, relative to their pangenome status. The color of each bar indicates the expression cluster to which lncRNAs are attributed. The numbers inside each bar show the raw counts of lncRNA candidates, while the y-axis shows the proportion of all candidates within each category (accessory or core) that belong to the cluster.

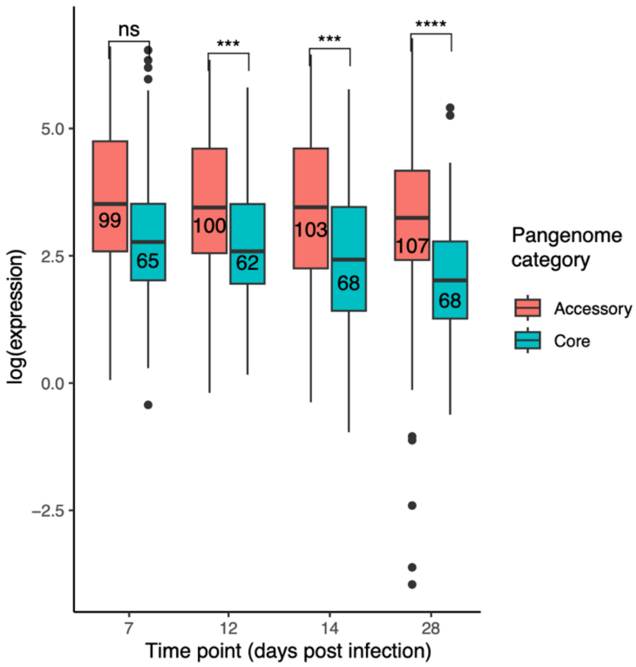


**Supplementary figure 16:** Box-plot comparing log TMM values between core (blue) and accessory (red) lncRNAs in four additional reference strains across the infection cycle. The numbers inside the boxes represent the sample size. The stars above each group of boxes shows the significance level of a welch two sided t-test comparing the two groups (indicated by the parentheses) at each time-point. Each time-point was considered independent and tested separately. The value under the parentheses shows the effect size of the difference between the two means.

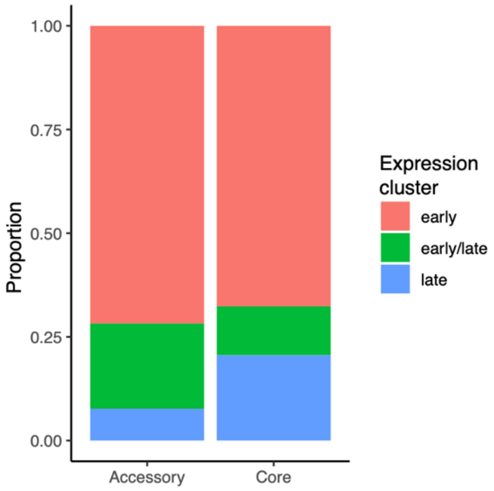


**Supplementary figure 17:** Bar chart showing the distribution of core and accessory lncRNAs into expression clusters grouped by peak expression. The y-axis shows the proportion of lncRNAs in each category (core or accessory) that are attributed to each group of expression clusters (peak expression is either early, late, or bi-modal).
